# Genetic and environmental imprints on T cell receptor repertoires as predictors of graft-versus-host disease

**DOI:** 10.64898/2025.12.23.696252

**Authors:** Assya Trofimov, Zachary Montague, Magdalena L. Russell, Rachel Bender Ignacio, Terry Stevens-Ayers, Danniel Zamora, Marco Mielcarek, Michael J. Boeckh, Frederick Matsen, Armita Nourmohammad

## Abstract

Hematopoietic cell transplantation (HCT) as potentially curative treatment for patients with hematologic malignancies relies on T cells to mediate the potentially curative graft-versus-tumor (GVT) effect, which may be associated with graft-versus-host disease (GVHD), a potentially life-threatening complication. T cells recognize peptides presented by Human Leukocyte Antigen (HLA) molecules. It is commonly believed that matching HLA alleles between donors and recipients ensures similarity in their T cell receptor (TCR) repertoires, thereby reducing the risk of GVHD and graft rejection. However, TCR repertoires are shaped by multiple factors beyond HLA genetics, including sex, age, and immune history. The extent to which genetic variation and past infections influence TCR repertoire composition and GVHD risk remains unclear.

Here, we show that while HLA haplotypes contribute to broad TCR repertoire differences, recent viral infections significantly impact TCR composition and influence GVHD risk. Analyzing 401 patients who were uniformly transplanted from healthy HLA-identical sibling HCT donors, we introduced *HLA-TCR coherence*, a metric that quantifies the extent to which an individual’s TCR repertoire reflects their HLA haplotype. We find that higher TCR-HLA coherence is associated with greater HLA heterozygosity and an increased incidence of severe acute GVHD (grade 3-4) in transplant recipients. Furthermore, *in silico* identification of virus-associated TCRs (vaTCRs) using TCR sequencing and viral serology reveal specific vaTCRs predictive of either increased or decreased GVHD risk. These findings suggest that beyond HLA allele matching, donor-specific immune history and repertoire characteristics are critical determinants of GVHD risk. Thus, a more systematic integration of donor immune history may offer a complementary avenue for refining donor selection and potentially improving transplant outcomes.

**One Sentence Summary:** While genetics shape the potential of a donor’s T cell repertoire, factors like infections and HLA diversity influence its composition and impact on transplant outcomes, including GVHD risk.

## INTRODUCTION

T cells play a key role in the adaptive immune system through surveillance and response to pathogens. To mount a response, T-cell receptors (TCRs) recognize short pathogen-derived protein fragments (peptides) presented on Human Leukocyte Antigen (HLA) molecules expressed on the surface of infected cells and antigen-presenting cells. To mount specific responses against the vast array of peptide antigens, our immune system generates a diverse repertoire of TCRs through a stochastic process known as V(D)J recombination, involving the rearrangement of variable (V), diversity (D), and joining (J) gene segments, along with random nucleotide insertions and deletions at junctions. Viral infections drive the expansion of T-cell clones that recognize viral antigens, enhancing the immune system’s ability to control and eliminate the infection. Consequently, an individual’s TCR repertoire reflects their history of pathogen exposure. In parallel, inherited HLA alleles play a critical role in determining which antigens are presented to the immune system and thus influence the composition of the responding TCRs. Despite recognition of these influences, the relative contribution of genetic factors (e.g. HLA alleles) versus environmental factors (e.g. pathogen exposure) in shaping TCR repertoires remains poorly understood. Disentangling this interplay is essential for advancing our understanding of immune adaptation and its implications for an individual’s susceptibility to immune-mediated diseases.

Current experimental approaches for probing for antigen-specific TCRs are prohibitively expensive. Unlike antibodies, which can recognize antigens in their native conformations, T cells require antigen presentation via HLA molecules. This necessitates the production of specific tetramers for each antigen under investigation, significantly increasing cost and complexity. Furthermore, even when T cells binding to antigen-HLA tetramers are identified, the results can be confounded by non-specific binding and false positives (Nagorsen *et al*., 2002; Newell *et al*., 2013). As an alternative, some studies have leveraged computational methods to infer antigen-specific TCRs. For instance, (Emerson *et al*., 2017) used machine learning to identify cytomegalovirus (CMV)-associated TCRs from repertoires annotated as CMV-positive or CMV-negative.

In this study, we leverage VirScan—a high-throughput serological profiling technique—to infer individuals’ viral exposure histories based on antibody reactivity to a panel of over 300 viruses. Notably, we use VirScan data from the same cohort for which TCR repertoires are available from (Emerson *et al*., 2017). While VirScan does not provide information about the timing of infection, it shows the relative abundance of antibodies reactive to specific viral peptides, which indicates either prior viral exposures or immunization. Using these data, we identify virus-associated TCRs and examine their relationship with each individual’s HLA alleles. Furthermore, we introduce a novel metric, *HLA-TCR coherence*, which quantifies the how much an individual’s TCR repertoire is described by their HLA haplotype. We show that high HLA-TCR coherence in hematopoietic cell transplantation donors is associated with an increased risk of graft-versus-host disease (GVHD) in recipients following transplantation. This suggests that while a coherent HLA-TCR relationship may enhance the immune system’s ability to target specific antigens, it may also narrow the tolerance to the minute inter-individual differences that are typically associated with GVHD. These findings highlight the interplay between genetic and environmental factors in shaping individual TCR repertoires and underscore their implications for immune-related diseases.

## RESULTS

### Experimental setup and feature engineering

The precise composition of the TCR repertoire is heavily dependent on an individual’s HLA haplotype (Ashby and Hogquist, 2024; Ashton-Rickardt *et al*., 1993; Brown *et al*., 2024; Johnson *et al*., 2021). Our recent study demonstrated that both age and sex as well as human CMV seropositivity also influence the TCR repertoire composition in a major way (Trofimov *et al*., 2022) (Figure 1A). Recently, a modest correlation between HLA overlap and TCR overlap was reported when comparing the occurrence of specific TCRs between individuals with similar or dissimilar HLA haplotypes (Krishna *et al*., 2020). However, due to TCR repertoires being highly undersampled (see details in Mora and Walczak, 2019), these results may not fully capture the relationship between HLA and TCR sharing. We hypothesize that a better signal between TCR repertoire similarity and HLA haplotype similarity can be seen at the level of general repertoire features such as V, D and J gene usage, CDR3β length and amino-acid position statistics. To verify this, for each individual in the *Emerson* cohort (Emerson *et al*., 2017), we trained a selection model (Isacchini *et al*., 2020) to characterize preferences for different TCR sequence features shaping individuals’ repertoires within this cohort (See Methods and Figure 1B). For better generalization of our predictions to unseen HLA haplotypes in our machine learning model evaluations, we grouped HLA Class I and Class II haplotypes by HLA binding cleft similarity and peptide binding preference (Thomsen *et al*., 2013) (see Methods) (Figure S1, Table S2). Finally, for information on individual immune histories, we performed VirScan (Xu *et al*., 2015) profiling of the same donor cohort (Manuscript in preparation). Taken together, these data offer a multi-modal characterization of the individuals in the Emerson cohort in terms of genetic and environmental features influencing their TCR repertoire composition.

**Fig. 1.**
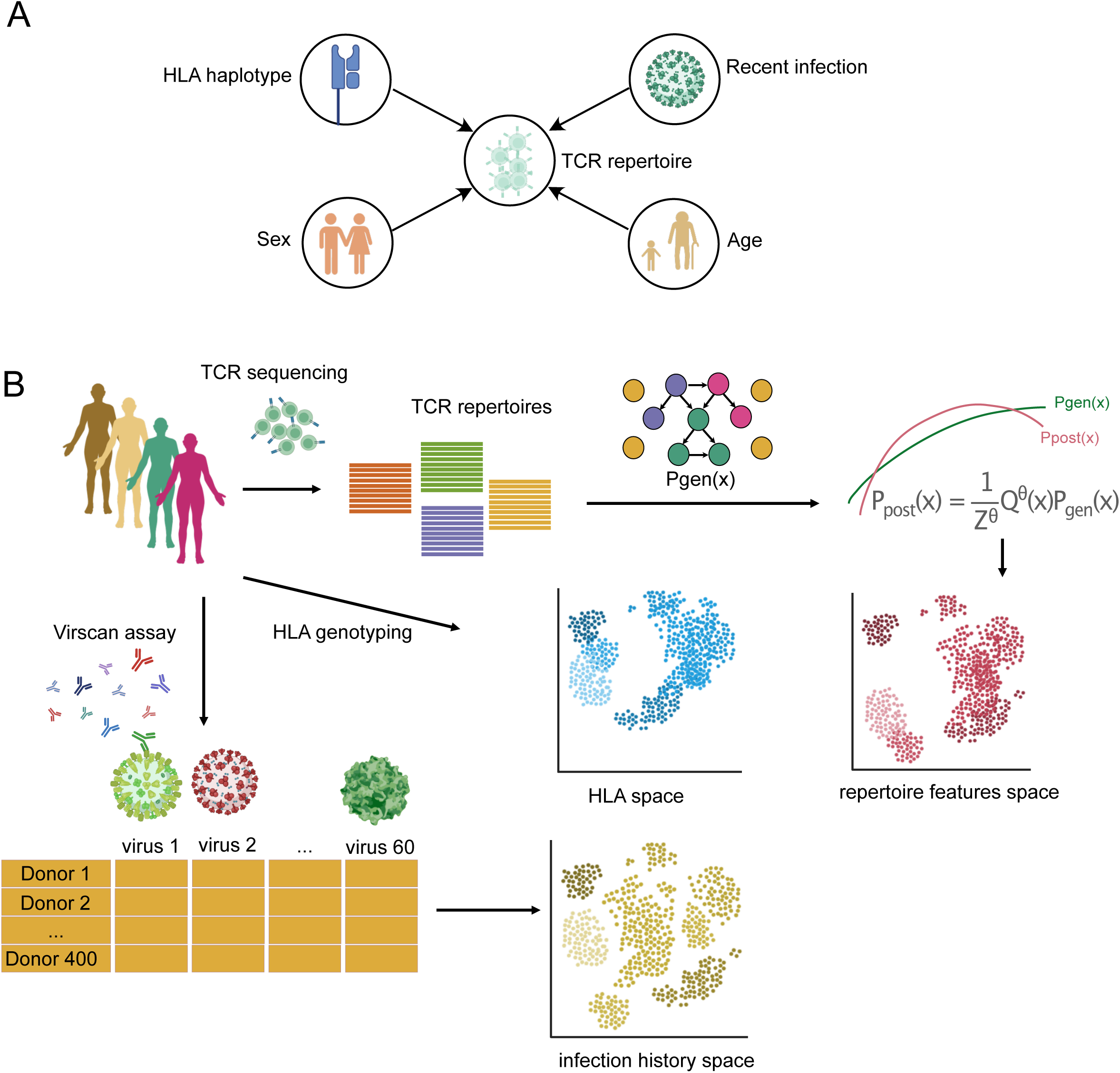
Experimental setup. **(A)** Schematized overview of examples of genetic and environmental elements that contribute directly to the composition of the TCR repertoire. **(B)** Schema showing data processing and experimental setup. For each TCR repertoire, a repertoire selection model (Isacchini *et al*., 2021) is trained (see Methods). For each individual’s HLA haplotype, HLA are encoded into the HLA space by clustering peptide binding clefts. VirScan assay readouts show a single value per virus per individual.

### Predicting TCR repertoire features from HLA haplotypes and infection history

To assess the relative contribution of HLA haplotype and infection history (via VirScan) to the TCR repertoire, we trained a contrastive neural network (see Methods) that predicts TCR repertoire *feature differences between individuals* from the differences in their HLA allele compositions, infection histories or a mix of both (Figure 2A; see Methods on details of model architecture). Next, we assessed prediction accuracy either on a *per-donor* basis or on a *per-feature* basis (Figure 2A, bottom right). Upon testing our model on a held-out set of individual repertoires (test set, see Materials and Methods for details), we found that in both types of comparisons, combining HLA with infection history did not yield a better performance than only using HLA (Figure 2B-C). Moreover, since viruses of the *herpes* family have been reported to have profound idiosyncratic effects on the composition of the TCR repertoire independent of HLA haplotype (Attaf *et al*., 2018; Emerson *et al*., 2017; Schober *et al*., 2020), we expected their corresponding VirScan features to have low predictive value. Herpesviruses such as CMV, EBV, and HSV are known to establish lifelong latency and can dominate the memory T cell compartment through large, individual-specific clonal expansions that do not generalize well across individuals—even those with similar HLA haplotypes (Jamaleddine *et al*., 2024). These persistent, idiosyncratic responses likely introduce substantial variability that obscures more generalizable repertoire features. Indeed, after excluding the herpesviruses from the inputs to the model, as expected, we found the performance dropped significantly (Tukey post-hoc p-value = 6.6x10^-11^) by 0.38, from a Pearson’s r=0.62 to Pearson’s r=0.24 (Figure 2B-C). Moreover, upon further investigation, we found that HLA data combined with VirScan data–but omitting the *herpesvirus* family from the inputs—yields the best accuracy of prediction (Pearson’s r =0.653) (Figure 2B-C). Finally, solely using viral infection history (without *herpesviruses*) resulted in a higher performance (Pearson’s r=0.621) than HLA only (Pearson’s r=0.585) (Tukey post-hoc p-value = 6.89 x 10^-8^) (Figure 2B-C). These results highlight the fact that both HLA haplotypes and viral immune history contain enough signal to make predictions of similar precision about general TCR repertoire features.

**Fig. 2.**
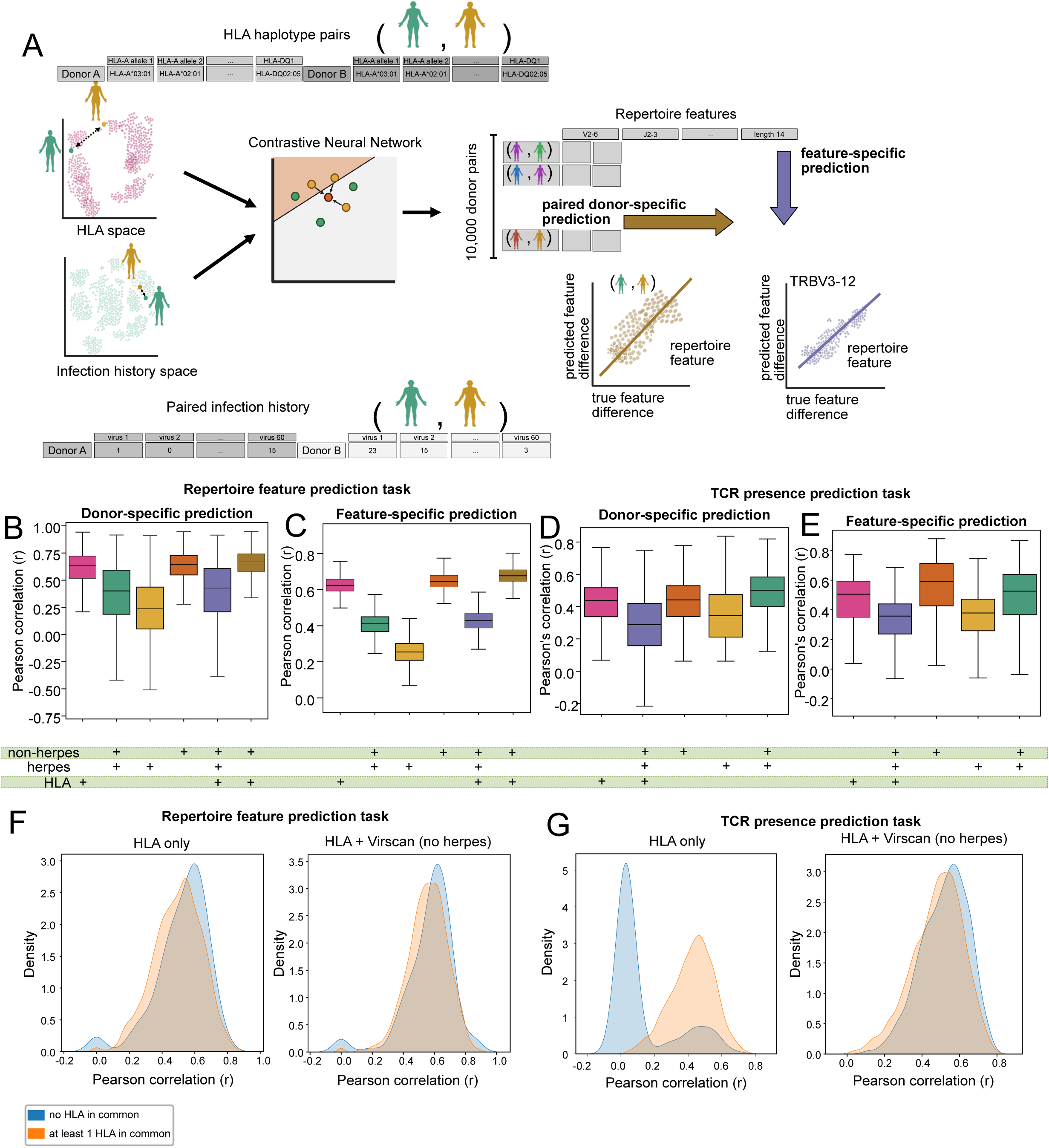
Predicting TCR repertoire features from HLA haplotypes and infection history. **(A)** Schematic of the contrastive learning setup. For every pair of individuals, HLA and/or VirScan data are input to a contrastive neural network model (see Methods). The model predicts repertoire features learned by repertoire models (Isacchini *et al*., 2021) (Figure 1B, Methods). Model performance is evaluated in a paired-donor- or feature-specific fashion by calculating Pearson correlation coefficient (r). **(B to D)** Boxplots showing the performance of the contrastive neural network on (B) TCR repertoire feature prediction task for various input modalities evaluated by paired-donor prediction and (C) feature-wise predictions measured by a Pearson correlation coefficient (r). Boxplots showing the performance of the contrastive neural network on TCR presence/absence prediction task measured by Pearson correlation coefficient (r). The model performances on each modality was evaluated by (D) paired-donor prediction and **(E)** feature-wise predictions. The bottom row indicates which dataset modalities or subsets were included in training the contrastive neural network. For all four sets of boxplots, each boxplot represents the performance for 10,000 data points and statistical significance was assessed by U-test and all comparisons were found to be statistically significant after multiple testing correction (corrected p-value <0.05). **(F and G)** Comparing HLA only and HLA+Virscan (no herpesvirus) as inputs, the performances (Pearson correlation coefficient (r)) of the contrastive neural network (F) on the repertoire features prediction task and (G) TCR presence/absence task are reported. Donor-pairs are split by number of HLA alleles in common, and the model performance distributions are displayed as KDE-smoothed curves.

We then trained a contrastive neural network model with different input combinations (HLA and VirScan information) to predict the presence or absence of randomly selected TCRs in a cohort subset. Specifically, we selected three sets of 1,000 random TCRs found in (i) 2-50, (ii) 50-300, and (iii) more than 300 individuals from our cohort (see Methods), and trained the model to predict TCR presence or absence in each pair of repertoires. This task parallels the benchmarking procedure described in (Krishna *et al*., 2020). For the TCR presence/absence prediction task, the hierarchy of data input performances was similar to the one found for the TCR repertoire model feature prediction task. We found that the best performance was achieved when combining HLA and VirScan without *herpesviruses* (Pearson’s r=0.487) (Figure 2D-E). While other performance comparisons all were statistically significant (adjusted p-value <0.05), the difference between using HLA only (Pearson’s r=0.443) and using VirScan without *herpesviruses* (Pearson’s r=0.445) was not statistically significant (adj. p-value = 0.73). Finally, we found that in the TCR-presence prediction task, most pairs of individuals that share no HLA alleles in common showed near-zero Pearson correlation (Figure 2F-G). This result indicates that HLA alone is insufficient for predicting the presence of specific TCR clones (Figure 2G), even though it may still capture broader repertoire features (Figure 2F). Taken together, these findings highlight that factors beyond HLA, most notably infection history, play a critical role in shaping the composition of an individual’s TCR repertoire.

### Disentangling the predictive signal from HLA alleles and recent immune challenges

To disentangle the relative contribution of individual HLA alleles to the overall prediction of repertoire features, we iteratively masked each set of HLA alleles from the input data to the model and compare the changes in overall predictive performance (see Methods). Unsurprisingly, we found that the prediction performance improves when more HLA alleles are presented to the model (Figure 3A). Using a linear regression, we modeled the relative contribution of each HLA allele to the overall performance. We found that when the model is provided with only a subset of the six alleles, specifically any combination of 1, 2, or 3 alleles, inputs that include either HLA-A or HLA-DRB1 consistently yield the highest prediction accuracy. This highlights the disproportionate influence of these loci on TCR repertoire features (Figure 3B–C). A similar conclusion was reached by examining SHapley Additive exPlanations (SHAP) values (Lundberg and Lee, 2017) for each allele type (Figure 3D). Our results are in line with those of Ishigaki et al., who have reported a strong association between HLA-DRB1 and the variability within the CDR3β region of the TCRs in the context of autoimmune pathogenicity (Ishigaki *et al*., 2022). Moreover, Walser-Kuntz et al. have also reported a strong association between the HLA-DRB1 and Vbeta gene usage in naive CD4 T cells. Walser-Kuntz and colleagues have found a less pronounced effect in memory CD4 T cells, underscoring the important effect of immune challenges in shaping the TCR repertoire composition (Walser-Kuntz *et al*., 1995).

**Fig. 3.**
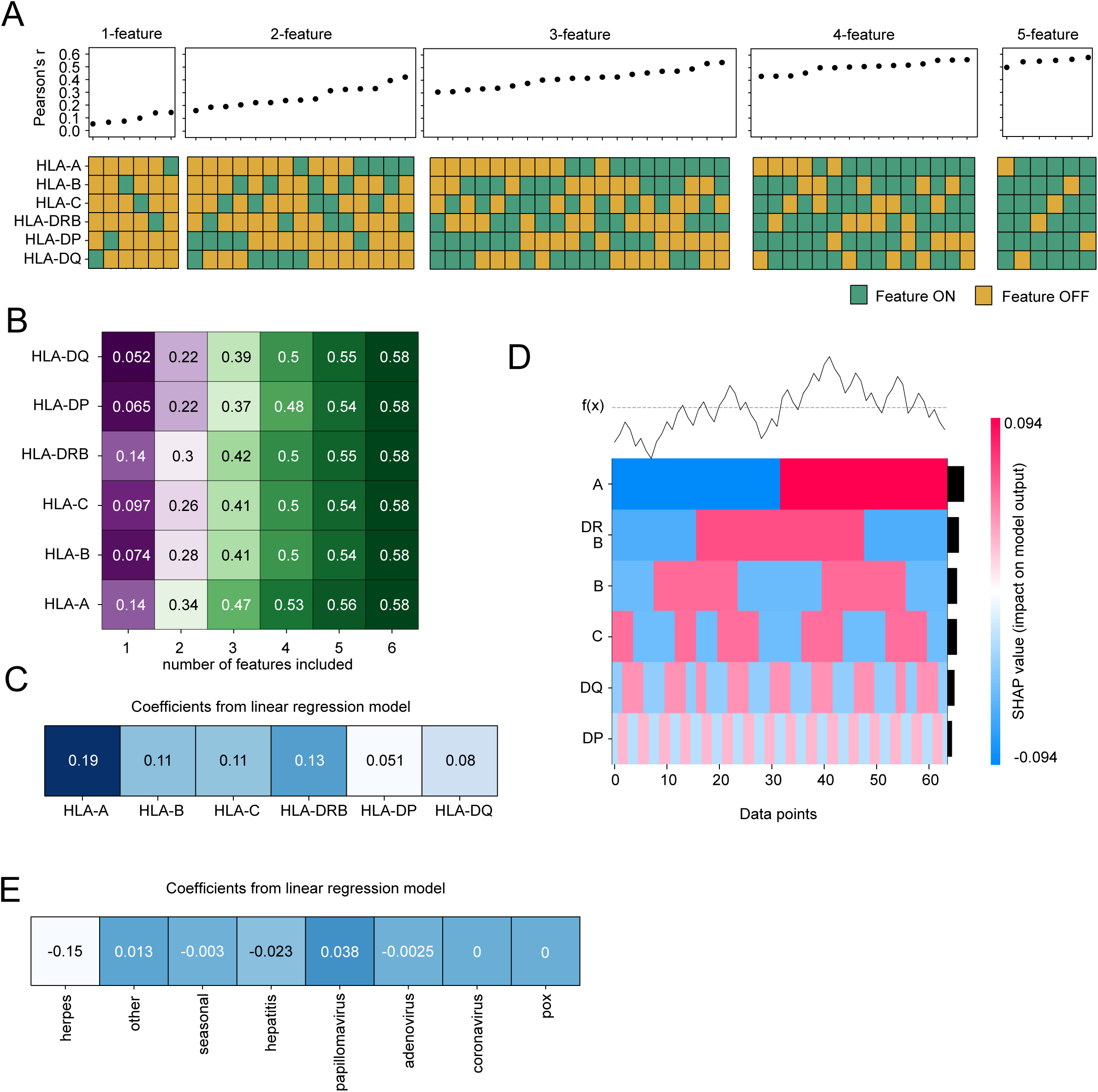
Disentangling the predictive signal for HLA alleles and recent immune challenges. **(A)** Performance of the contrastive neural network on the TCR repertoire feature prediction task as a function of HLA alleles presence Inputs are grouped by the number of alleles presented to the model, and performance measured with Pearson correlation coefficient (r) is shown in ascending order. The inclusion or exclusion of an allele is indicated according to the color-scheme in legend. **(B)** Mean performance (Pearson correlation coefficient r) grouped by number of alleles presented to the contrastive neural network and specific allele presence. Each square in the heatmap represents the averaged value for each feature number block from (A). **(C)** A linear repression model was trained to predict model performance (Pearson correlation coefficient (r)) from A from the different allele inclusion/exclusion binary vectors (heatmap in A). The coefficients from the linear regression model (arbitrary units) are shown as a heatmap.**(D)** SHAP values (Lundberg and Lee, 2017) based on individual allele contribution to TCR feature prediction task performance are visualized as a heatmap showing the total impact on the model’s performance. **(E)** The procedure from A-C was repeated for virus groups (*plot not shown)*. Coefficients from the linear regression model (arbitrary units) trained on iteratively masked VirScan input data are shown as a heatmap.

In a similar fashion, we grouped the viruses from the infection history data into categories based on families or infection timeline (Table S3) and found that responses to *herpesviruses* have a negative effect on the prediction overall, as we have observed previously (Figure 3E). Taken together our results show a heterogeneous contribution of HLA alleles and recent infections to the overall TCR repertoire composition, with HLA-A and HLA-DRB1 holding the most predictive power and responses to herpesviruses being the least predictive.

### HLA-TCR repertoire coherence as a measure for predicting GVHD

In the context of hematopoietic cell transplantation (HCT), it is assumed that a complete HLA match between a donor and a recipient will ensure the graft T cell repertoire to be closest to the recipient’s (Figure 4A). Indeed, this potentially life-saving treatment has a substantial risk of acute graft-versus-host disease (aGVHD), a potentially deadly complication of HCT. Conversely, some HLA mismatches are found to be more permissive (Fleischhauer *et al*., 2012; Martin, 1991), further highlighting the unpredictable nature of the disease. Moreover, it is generally accepted that minor histocompatibility antigens drive aGVHD in HLA-matched donor and recipient pairs(Baron *et al*., 2007; Martin, 1991; Summers *et al*., 2020). Currently, the relative contribution of HLA versus immune history, age and sex to the aGVHD risk is not known (Figure 4B). To better clarify these contributions, we leveraged the Emerson cohort, analyzing a total of 286 HLA-identical sibling donors as well as 115 mismatches or HLA-matched but unrelated donors (Table S1) (Emerson *et al*., 2017). We quantified the distance between TCR repertoires of all donor/recipient pairs in the Emerson cohort, using the Jensen-Shannon divergence (*DJS*) between selection models trained separately on each individual’s TCR repertoire (see Methods for details) (Figure 4C); high DJS values signify divergent TCR repertoires, while low values indicate similarity. As expected, DJS generally decreased as donors shared more HLA alleles. However, numerous donor pairs with no shared alleles still showed lower DJS values than some pairs with some allele overlap (Figure 4C). This result suggests that TCR repertoires sharing similarities can emerge even in the absence of HLA overlap.

**Fig. 4.**
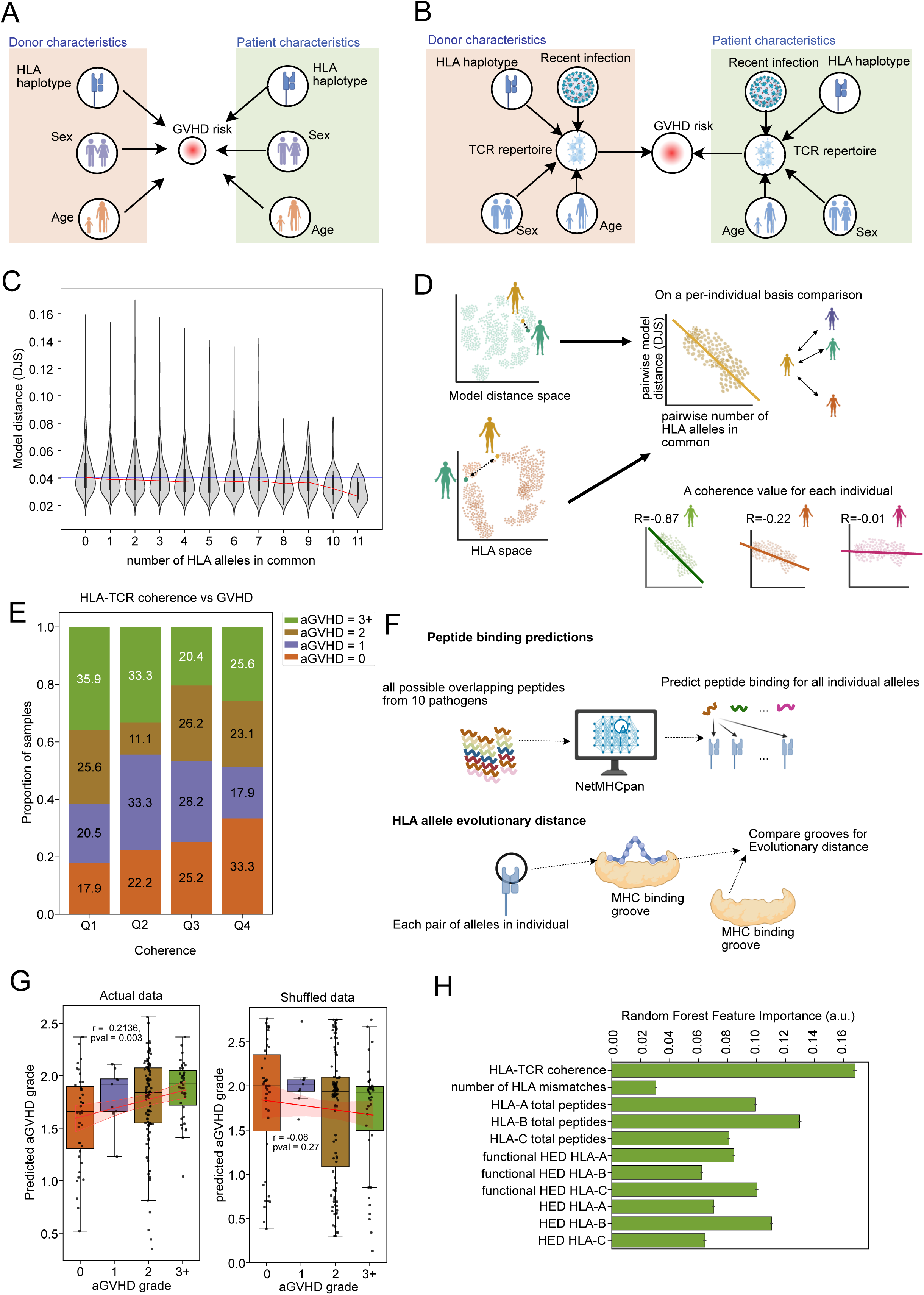
HLA-TCR repertoire coherence as a measure for predicting GVHD. **(A)** Graphical schematic illustrating conventional aGVHD risk assessment based on donor and recipient HLA, sex and age compatibility. (B) Updated risk evaluation framework incorporating both T cell receptor (TCR) repertoire features and immune history, highlighting missing TCR repertoire node. **(C)** For each TCR repertoire in our cohort, a selection model (Isacchini *et al*., 2021) was trained. Pairwise Jensen-Shannon divergences (DJS) as a measure of selection model distances was calculated for all pairs of individuals in the cohort. Violin plots show the pairwise model distance stratified by pairwise number of alleles in common for all pairs of individuals in the cohort. Blue horizontal line shows the median model distance for pairs of individuals with no alleles in common. Red line shows the median model distance for each number of of HLA in common **(D)** Schematic representation of the HLA-TCR coherence calculation. For each individual in the cohort, all pairwise model distances and all numbers of HLA in common are calculated. The HLA-TCR coherence is the Pearson correlation coefficient (r) calculated between these two measures of distance. **(E)** Individuals are binned into quantiles (Q1 = 0.25, Q2 = 0.5, Q3 = 0.75), Proportions of aGVHD severity grades is shown for each HLA-TCR coherence quantile. **(F)** Schematic representation of the HLA-based feature calculations: 1) for all possible 9-12 mers from 10 viral proteomes (Supplementary Table 2) peptide biding predictions are assessed using NetMHCpan(Reynisson *et al*., 2020). Strong binders are caracterized by a 0.5% binding ratio threshold. For individual’s MHC Class I allele, the size of the putative peptide pool is reported. 2) For each pair of MHC Class I alleles in each individual, the pairwise HLA allele evolutionary divergence (Radwan *et al*., 2020) is calculated by scoring each amino-acid position of MHC Class I binding cleft using Grantham distance (Grantham, 1974). 3) functional HLA allele divergence is calculated as a measure of overlap (Jaccard Index) between pairs of MHC Class I alleles using the peptide binding predictions from (1) **(G)** Random forest regressor models are trained to predict aGVHD grade severity from HLA-based features such as total predicted peptides and HLA allele evolutionary divergence from (F) as well as HLA-TCR coherence and number of HLA mismatches between the donor and patient. Their performance is assessed with a Pearson correlation coefficient between the predicted and actual aGVHD severity grade **(H)** Overall feature importances for the Random Forests from (G) are reported as a barplot..

We wondered if this repertoire similarity in the absence of HLA overlap was individual-specific. If so, we would expect some inter-individual variability, with certain donors showing stronger alignment between HLA and TCR distances than others. To test this, we compared each individual’s HLA haplotype and TCR repertoire model to those of all others in the cohort. Specifically, for each pair of individuals we obtained an HLA allele overlap in number of alleles (*HLA haplotype distance*) and the DJS between the two TCR repertoire models (*repertoire distance*). Next, we compiled all the pairs of distances between a single individual and the rest of the individuals in the cohort and computed the Pearson coefficient of correlation between these distances. The obtained Pearson correlation coefficient is this individual’s *HLA-TCR coherence*, which captures the extent to which differences in an individual’s TCR repertoire align with differences in HLA haplotypes across the cohort. A strongly negative correlation indicates a highly *HLA–TCR coherent* individual—that is, for that person, individuals who are more dissimilar in HLA also tend to be more distant in TCR repertoire. In contrast, correlations near zero indicate HLA–TCR incoherence, meaning that for that individual, HLA similarity (or dissimilarity) shows little alignment with TCR repertoire similarity (Figure 4D).

Splitting our *HLA-TCR coherence* measure distribution into quartiles, we found that individuals with the highest coherence (Q1) had the highest proportion of severe aGVHD (grade II-III) (Figure 4E, green box), whereas those with the lowest coherence (Q4) had a greater proportion of mild or no aGVHD (grade 0–1) (Figure 4E orange box). Next, we put together a list of general HLA features (Figure 4F), such as predicted peptide binding pool (see Materials and Methods, Figures S2) as well as HLA Evolutionary Distance (HED) (Di *et al*., 2020) between alleles and the HLA-TCR coherence, to assess the predictive potential of HLA-centric features for aGVHD severity grade. We trained a Random Forest regressor to predict aGVHD grade using these features, and our model demonstrated a modest but significant positive correlation (Pearson’s r = 0.2136, p-value = 0.003) between predicted and actual aGVHD grades on a held-out subset of samples (Figure 4G). A shuffled dataset resulted in a non-significant correlation (Pearson’s r = -0.08, p-value = 0.27), confirming the model’s validity (Figure 4G). Importantly, coherence emerged as the most influential feature in the model, surpassing other classical HLA-based metrics in the feature importance analysis (Figure 4H). Overall, these findings indicate that HLA-TCR repertoire coherence shows a modest association with aGVHD severity and may provide additional prognostic value beyond traditional HLA-matching approaches for aGVHD risk estimation.

### Infection history of the donor influences risk of acute graft-versus-host disease

Next, we sought to assess the link between infection history and occurrence of aGVHD after transplant in our cohort. We hypothesized that some transplants may be riskier because of the breadth of the donor’s viral exposure history and its potential imprint on the TCR repertoire. Alternatively, the history of infection may be associated with a proinflammatory milieu in the recipient, which could promote donor T cell activation and GVHD. Furthermore, infections in the recipient in the early post-transplant period can promote circulating donor T cell activation and increase the risk of GVHD. We assessed the association between the VirScan profile and the development of severe aGVHD using a Cox proportional hazards model (CoxPH) compared to clinical data variables and known predictors of aGVHD such as donor age, sex mismatch, CMV serostatus pairings, GVHD conditioning intensity and prophylaxis regimens. We found that sex mismatch or CMV serostatus pairing between the donor and patient did not have a significant effect on aGVHD risk and while having a large age gap (>50%) between the donor and the recipient was not statistically significant (corrected p-value = 0.32). Having an unrelated donor was associated with a 5.78-fold increase in hazard (corrected p-value = 0.002). In addition, higher HLA–TCR coherence was associated with an increased hazard ratio of 1.86 (corrected p-value = 0.003), in line with our observations (see Figure 4). For the VirScan profile associations, we report that a history of infection associated with *Human herpesvirus 6B, Human coronavirus HKU1* and *Human adenovirus D* in the donor had a higher hazard ratio of unfavorable outcome for graft recipients, while infections with *Orf virus, Human adenovirus C* and *Herpesvirus 3* are protective (Figure 5A). A possible biological mechanism that would make an infection protective could be the infection triggering the expansion of virus-associated T cell clones, which in turn could be less alloreactive, once transplanted into the recipient (Barrett, 2011; Litjens *et al*., 2018). Moreover, while overall viral exposure is a surrogate for donor age, where older individuals have had more time for more viral exposure (Bender Ignacio *et al*., 2019) (Figure S3A), this measure did not hold for the viruses our model found to be associated to aGVHD risk (Figure S3B). We verified these associations with single virus Kaplan-Meier curves split on the median for the VirScan profile and found statistically significant log-rank associations only for *Orf virus*, *Human herpesvirus 6B* and *Human coronavirus HKU1* (Figure S4A,B,D top plots). We also plotted single-virus KM curves for 6 randomly selected control viruses and found some statistically significant associations for *Molluscum contagiosum* and *Rhinovirus B* (Figure S5A,E top plots). However the CoxPH model by design does not split the samples into groups such as log-rank but rather evaluates risk as a function of time. The univariates CoxPH models were not statistically significant for the control viruses (bottom plots Figures S5) but were statistically significant associations for the viruses from Figure 5A (Figure S4, bottom plots). These two comparisons (univariate CoxPH and KM curve + log-rank) underline the need to use multivariate CoxPH, which adjusts for variable other variables such as clinical data, preparatory regimen and prophylaxis.

**Fig. 5.**
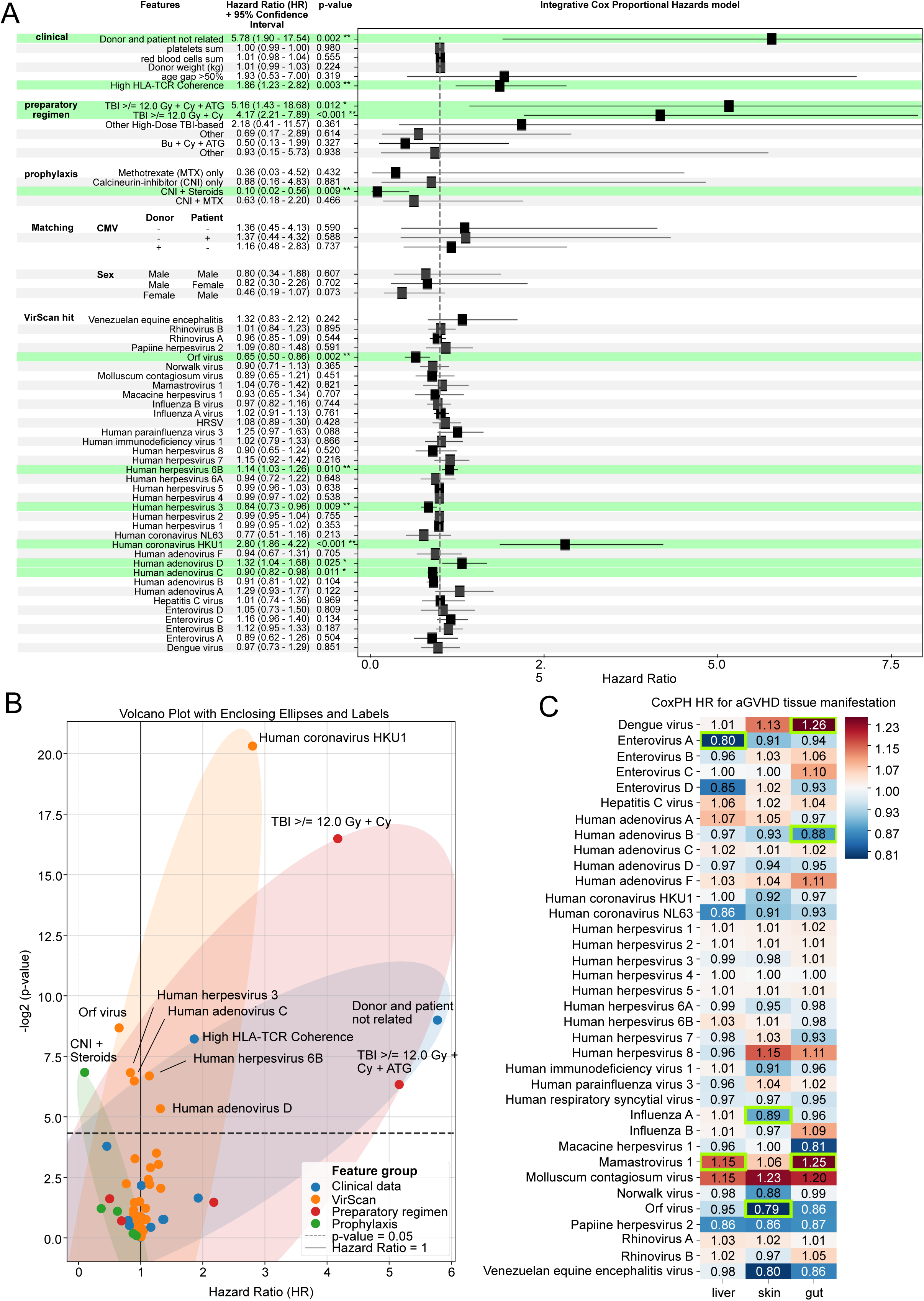
Modeling aGVHD occurrence from VirScan with survival models. (A) Results of the Cox Proportional-Hazards (CoxPH) model evaluating the association of each clinical variable, preparative regimen, prophylaxis, and virus-associated VirScan signal with time to onset of aGVHD, For each feature, the model’s calculated hazard ratio (HR) and 95% confidence interval (CI) are shown as well as corrected p-value. Predictive variables with corrected p-value <0.05 are highlighted in green **(B)** Scatterplot representation of the CoxPH results, showing hazard ratios (HR) plotted against -log2(corrected p-value) for all clinical variables, preparative regimens, prophylaxis and viruses. Gaussian kernel density ellipses outline the broad distributions for each feature category, providing a visual summary of group HR and significance trends. **(C)** CoxPH model HR results stratified by aGVHD tissue manifestations shown for each virus. Green squares signify a significant corrected p-value for that virus. Values are coloured in blue for lower risk features (HR <1) and red for higher risk (HR>1).

While all aGVHD prophylaxis regimens had a protective effect (general HR values <1), preparatory regimens with total body irradiation (TBI) all had hazard ratios for aGVHD occurrence higher than 1 (Figure 5A-B). A caveat is that we did not evaluate relapse risk in the non-TBI regimens, and determining whether this contributes to better graft outcome overall will require additional study. In general, the hazard ratios calculated by the CoxPH for viral exposure were centered around 1 and had a smaller spread (and effect) compared to those of the prophylaxis and preparatory regimens (Figure 5B). Finally, to assess potential links between viral tissue tropism and aGVHD tissue manifestations, we repeated the CoxPH model calculation this time predicting tissue manifestations of aGVHD in the lung, skin and gut from the VirScan data. We found that for liver aGVHD, a donor infection history of *Enterovirus A* was protective (HR = 0.8) while *Mamatrovirus 1* was unfavorable (HR = 1.15). Similarly, we found that a history of *Influenza A* (HR= 0.89) and *Orf virus (*HR= 0.79) infection were protective for skin aGVHD manifestation, and *Human adenovirus B (*HR= 0.88) was protective for gut aGVHD manifestation, while *Dengue virus* (HR= 1.26) and *Mamastrovirus 1* (HR= 1.25) were harmful (Figure 5C). While for some viruses a link between the virus tissue tropism and the aGVHD association can be made (*Orf virus* and skin aGVHD, *Mamastrovirus 1 and Adenovirus B* and gut aGVHD), for others the link is not direct. It is important to note that a positive VirScan signal should be interpreted not as definitive seropositivity, but rather as a marker of an immune response linked to that specific virus (Xu *et al*., 2015). This nuance may help explain some of the indirect associations between viral tissue tropism and aGVHD manifestations, as well as positive VirScan hits for rare viruses in the general population (ex: *Orf virus, Cowpox virus, Dengue virus*). Taken together these data show that there is a statistically significant association between severe aGVHD risk and donor infection history but that more investigations in additional cohorts are needed.

### Identifying virus-associated TCRs *in silico*

To further investigate the associations between viral immune repertoire and the risk of developing aGVHD, we developed a pipeline to identify virus-associated TCRs (vaTCRs) using the VirScan and TCR repertoire data. We hypothesize that a donor’s viral exposure profile points to changes in the abundance or presence of certain virus-associated TCRs in the donor and that these TCRs could be predictive of aGVHD development. We use a series of three filters to refine vaTCRs from the 80 million unique TCRs in the dataset (Figure 6A) (see Methods for details). Briefly, we restricted the TCRs to public TCRs (seen in at least 2 individuals), reducing the set to about 2 million TCRs. Next, for each TCR, we trained a logistic regression model to predict the likelihood of observing a TCR in an individual, given the complete viral exposure or immunization history of that individual (from VirScan). We kept TCRs for which the predictive performance on a held-out set of individuals is higher than that of shuffled data and that difference was statistically significant (corrected p-value <= 0.05), further reducing the TCR set to about 10000 infection profile-associated TCRs (Figure 6A). Then, as a final filter, we performed a series of Fisher tests using a contingency matrix constructed on the co-occurrence of a TCR with a high VirScan readout for a single virus (corrected p-value <= 0.05, see Materials and Methods). This last filter associates each infection profile-associated TCR to a single virus to extract the final list of vaTCRs (Figure 6A). With this method, we isolated 4577 vaTCRs, with between 2 and 390 vaTCRs per virus (Figure S6A). Using multiple reshuffles on our data we calculated a false discovery rate of 4% for the entire enrichment procedure (see details in Methods, Figure S7A-B). The vast majority of vaTCRs were associated with a single virus but a handful were shared between two viruses. There was no clear pattern among viruses that shared TCR associations: viruses that had TCR associations in common were not in the same family and were not expected to share any epitopes, except for a single pair: *Molluscum contagiosum* and *Orf virus* (Figure S6B). We also confirmed that virus pairs sharing vaTCRs do not show increased co-occurrence in VirScan profiles. For each virus pair, the observed correlations were modest, and the overall distribution of these correlations was indistinguishable from that of randomly selected virus pairs (Figure S6B-C). Cross-referencing vaTCRs with VDJdb (Shugay *et al*., 2018) and McPAS (Tickotsky *et al*., 2017) revealed minimal overlap, with only two matching associations: CASSPGTLYGYTF (Dengue) in VDJdb and CASREGGNQPQHF (HKU1/SARS-Cov-2) in McPAS, indicating a modest shared virus specificity. We also found 8 vaTCRs associated to *Human coronavirus NL16* and *HKU1* in our pipeline in the published 160,000 high-confidence SARS-CoV-2-specific TCRs from Adaptive Biotech’s ImmuneCODE (https://doi.org/10.21417/ADPT2020COVID). Although these databases are far from comprehensive, even a modest overlap helps validate our pipeline, especially since the overlap between our coronavirus-vaTCRs and these resources is similar to the overlap observed among McPAS, VDJdb, and MIRA (Figure S6D). Next we assessed the sequence features of the vaTCRs and report that the vaTCRs shared between viruses were found to have a shorter CDR3β length in amino acids (Figure 6B) and a higher probability of generation under a general V(D)J recombination model (Figure 6C), pointing towards their identity as public TCRs occurring at high prevalence in the population (Elhanati *et al*., 2018; Pogorelyy *et al*., 2017).

**Fig. 6.**
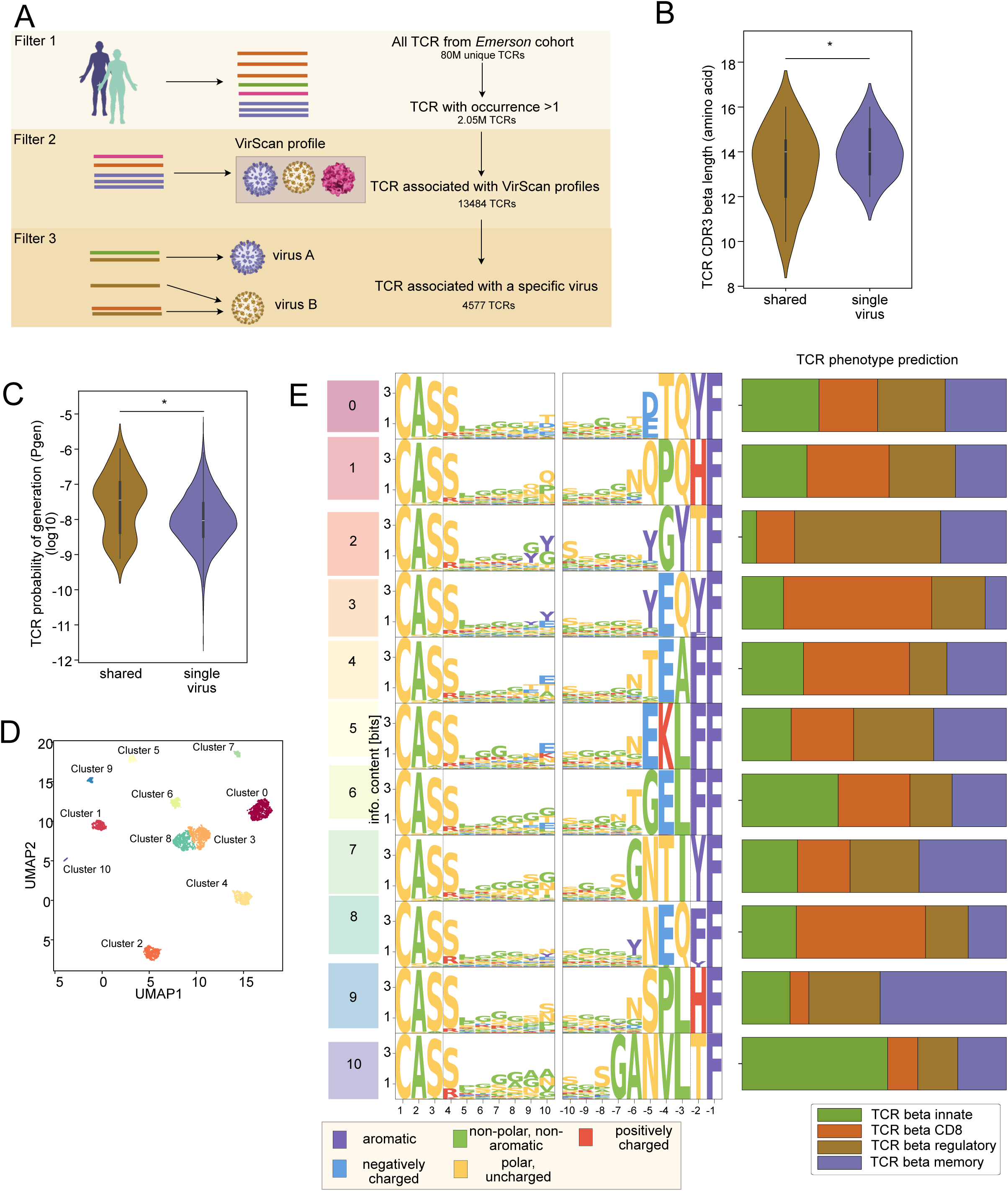
Computational pipeline to extract virus-associated TCRs. **(A)** Schematic representation of the multi-step enrichment pipeline (see Methods) for enrichment of virus-associated TCRs (vaTCRs). Starting from TCRs seen in at least 2 individuals in the cohort, logistic regression models are trained to predict presence or absence of a TCR given an individual’s VirScan profile. TCRs that show positive signal (AUC-ROC >0.6 and statistically different from randomly shuffled set) are retained for the next filter. Fisher tests are applied to find associations between a single virus and each retained TCR, only statistically significant associations (corrected p-value) are qualified as vaTCRs. **(B and C)** Violin plots of (B) TCR length distributions in amino-acid and **(C)** predicted generation probability (Pgen) (Sethna *et al*., 2019) for vaTCRs associated to a single virus and vaTCRs shared between viruses. Statistical significance is assessed by t-test, p-value<0.05. **(D)** Pairwise distances between all vaTCRs were calculated with *tcrdist* (Dash *et al*., 2017). K-Means clustering was performed on pairwise distance matrix to find vaTCRs clusters. Plot shows UMAP reduction of the pairwise vaTCRs distance matrix with vaTCRs coloured by cluster. **(E)** For each vaTCR cluster from (D), TCR phenotype proportions were predicted using *tcrpheno* (Lagattuta *et al*., 2024). TCR sequence logoplots (left) as well as TCR phenotype proportion predictions (right) are shown for each cluster from (D).

### Molecular patterns of vaTCRs

To get a sense of the molecular patterns of these vaTCRs, we used *tcrdist* (Dash *et al*., 2017; Mayer-Blackwell *et al*., 2021) to calculate a pairwise distance for each pair of vaTCRs and visualized this distance graph using uniform manifold approximation and projection (UMAP) (Figure 6D). With k-means clustering (see Methods), we obtained for each vaTCR their cluster membership based on their *tcrdist* measure. For each cluster, we visualized the logo plot (Figure 6F) and found distinct patterns in the different TCR clusters. We however did not find in these clusters any specific enrichment for hits from known peptide-TCR databases (Chronister *et al*., 2021; Shugay *et al*., 2018; Tickotsky *et al*., 2017) (Figure S8). We also used the recently published TCRpheno model that predicts TCR fates given TCR sequences (Lagattuta *et al*., 2024). We found that certain clusters were enriched in specific T cell compartments: clusters 3 and 8 were enriched in CD8 T cells (Figure 6E), cluster 10 was enriched in innate T cells, cluster 2 in regulatory T cells, and cluster 9 in memory T cells (Figure 6E). This finding aligns with well documented cross-reactivity issues in TCR datasets (Banerjee *et al*., 2024).

### Abundance of vaTCRs for certain viruses is predictive of aGVHD outcome

Next, we looked at the association between vaTCRs and the occurrence of aGVHD. We found that donors whose recipients developed aGVHD had a higher amount of vaTCRs deemed as “dangerous” from our survival analysis results (Figure 5A) detected in their repertoires (Figure 7A). In a similar fashion, for "protective" viruses, we found a negative but not statistically significant correlation between aGVHD severity and number of "protective" vaTCRs (Figure 7B). A random selection of viruses did not show any correlation (Figure 7C). Moreover, we found no association between aGVHD severity and the donor’s repertoire clonality (Figure 7C), highlighting that the association from Figure 7A-B is not simply a matter of TCR diversity but may depend on the details of repertoire composition, such as the presence of vaTCRs. Finally, relating this result to HLA-TCR coherence, we found that highly *coherent* individuals (Q1) had a greater number of vaTCRs (Figure 7D). Taken together these results show that the vaTCR abundance profile is associated with aGVHD severity, and this warrants further investigation to assess the contribution of VirScan data and vaTCR enrichment to aGVHD risk predictions (Figure 4B).

**Fig. 7.**
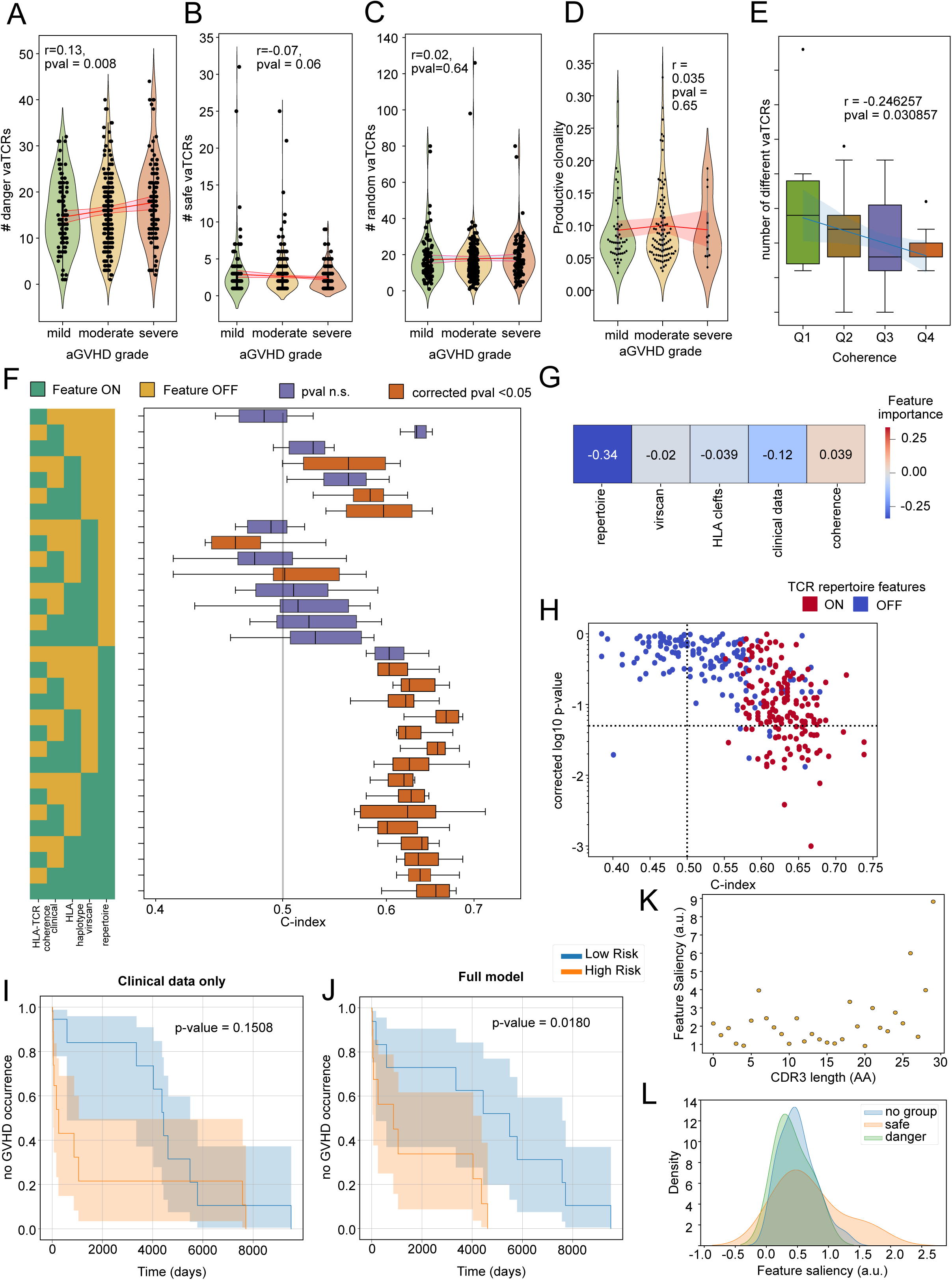
An integrative model underscores the importance of TCR repertoires in predicting GVHD risk. **(A and B)** Violin plots showing the numbers of vaTCRs present in each TCR repertoire in our cohort as a function of patient aGVHD severity. The vaTCRs are stratified by associated virus and split into **(A)** danger and **(B)** safe viruses from our previous results (Figure 5A). **(C)** Violin plot showing productive clonality of TCR repertoires by aGVHD severity. **(D)** Boxplot showing the number of different vaTCRs per sample, split by HLA-TCR coherence quartiles (Figure 4E). Pearson correlation coefficient (r) and associated p-value is indicated on each plot. **(E)** During the training of the Deep learning survival model, features were iteratively masked to assess the change in model performance based on their presence/absence. The heatmap on the left shows for each model which features were used (green = ON, yellow = OFF) and the boxplots show the corresponding model performance evaluated by C-index. Each boxplots represents 50 reshuffles of the data with a 80-20 split between training and test set. Boxplot color indicates statistical significance of log-rank test by splitting on the median the risks predicted by each model. **(F)** Similar to Figure 3, a linear regression model was trained to predict C-index from dataset on/off binary vectors from Figure 7E. Feature importance from this linear regression are reported as a heatmap. **(G)** Scatterplot showing C-index vs corrected log-rank log10 p-value for survival tasks with runs including (red) or exclusing (purple) TCR repertoire features data. **(H)** Kaplan-Meier plots are shown comparing the predicted aGVHD risk values split by median value for a deep survival model trained on clinical data only or **(I)** all data modalities (ex: TCR-HLA coherence, TCR reperotire features, VirScan, clinical variables) as inputs. Log-rank p-values are indicated for each plot. **(J)** Feature saliency map is shown for a deep survival model for TCR amino-acid lengths and **(K)** vaTCR associated virus “safe” or “danger” group membership.

### An integrative model underscores the importance of TCR repertoires in predicting aGVHD risk

Finally, in order to reconcile all the different genetic and environmental effects we studied that may influence the TCR repertoire, and in turn may contribute to aGVHD risk, we built a deep learning model to predict aGVHD disk from various genetic and environmental inputs. Our model is inspired by DeepSurv (Katzman *et al*., 2016) but uses discrete time transformers instead of fully connected layers (see Methods). Specifically, this artificial neural network receives as input any number of dataset modalities, including TCR repertoire features, clinical data as well as HLA features, and predicts a risk score. This risk score is a continuous scalar output from the neural network’s final layer, representing the log-risk of developing aGVHD for each patient. In DeepSurv and in our model, the risk score reflects a relative risk: individuals are compared against others who are still at risk at the same time *t*. The scale of the score is arbitrary, and only the ordering of risk scores affects the loss function. This ordering comes from comparing the times to the onset of aGVHD for each individual. Higher scores correspond to higher hazard and earlier expected onset of the event. These scores provide a relative ranking of patients’ risks, in line with the Cox proportional hazards assumption (Katzman *et al*., 2016). As previously described, in order to disentangle the relative importance of the various donor features, we iteratively masked inputs (Figure 7F). We evaluated the model using the concordance index (C-index), which measures how well individuals are ordered by predicted risk; a C-index of 0.5 corresponds to a random ordering while the C-index of 1.0 corresponds to a perfect ordering. We observed that models that included TCR repertoire features together as inputs (e.g. V,J gene frequencies, CDR3 lengths and amino-acid positioning) consistently had higher C-index values, compared to those with masked TCR repertoire features (Figure 7F-H). As for other data modalities such as clinical data, HLA-TCR coherence and VirScan , while their individual contribution to the model performance seemed low (Figure 7G), they had an additive importance, as can be seen in the stairs-like performance boost with added unmasked data modalities (Figure 7F). Indeed, the best performing model was the one that included all data modalities together (Figure 7F). It is important to note that while the model that only included clinical information such as sex, age and CMV serostatus of the donor and recipient performed well in terms of C-index, the calculated log-rank test was not statistically significant (Figure 7F,I), making this model inadequate at separating Low and High Risk individuals. In contrast, the model trained on the full selection of data modalities had both a C-index above 0.5 and a statistically significant p-value (Figure 7J). Next, we examined the saliency values over the input features to isolate the ones that contribute the most to the prediction. We found that when looking at CDR3β amino-acid length, TCR repertoires with distributions skewed towards longer length have a stronger contribution to model predictions (Figure 7K). In general, longer CDR3β amino-acid length is a hallmark of private, donor-specific sequences, seldom shared (Murugan *et al*., 2012). Also, the model attributed higher scores to Class I HLA and specifically in the diversity of peptides bound by HLA-A rather than Class II (Figure S9A). This interpretation should be made with caution, as peptide binding predictions for Class II HLA alleles remain considerably less accurate and more variable than Class I predictions (Backert and Kohlbacher, 2015). Consequently, the stronger attribution observed for Class I (and particularly HLA-A peptide diversity) may partly reflect differences in prediction reliability across HLA classes. In terms of specific alleles, HLA-B and HLA-DP cluster memberships were the most important, compared to other HLA (Figure S9A), which is congruent with previously outlined risk associations for HLA-B and HLA-DRB1 (Petersdorf *et al*., 2007). Other features were important, including V-J gene combinations as well as specific VirScan virus readouts (Figure S9B). Interestingly, the model attributed a higher importance to the VirScan assay inputs of protective viruses, rather than the risky ones or the ones without annotation (Figure 7L). Taken together, these results suggest that TCR repertoire data and related VirScan data may have relevance to aGVHD risk, warranting further exploration.

## DISCUSSION

In this study, we explored the interplay between genetic and environmental influences on shaping the human TCR repertoire, focusing on the relative contributions of HLA haplotypes and viral infection history. Using a large multi-modal dataset that includes TCR repertoire sequencing, HLA typing, and VirScan-derived infection profiles, we demonstrated that both the inherited HLA variation and the prior immune exposures exert measurable and complementary influences on the composition of TCR repertoires. Comparing selection models rather than overlaps in CDR3 sequences yields a coarse-grained repertoire-level comparison that is more robust to the catastrophic undersampling in TCR repertoires. More specifically, this type of comparison is able to account for the statistics of the missing sequences that have a likelihood of being seen in an individual if all the T cells were sampled. We were then able to more robustly measure the impact of an individual’s HLA composition and infection history on the model-derived statistics of their TCR repertoires. We found that while HLA by itself is predictive enough of TCR repertoire statistics, incorporating infection history data from VirScan slightly enhanced the predictive power. The VirScan-associated improvement was most pronounced when modeling the presence of specific TCRs instead of repertoire statistics. However, we found that not all viral exposure signatures contribute positively: VirScan data from herpesviruses consistently reduced model performance. This suggests that chronic herpesvirus infections might introduce noise into the analysis of TCR repertoires, rather than providing informative signatures that are predictive of overall repertoire composition. One likely mechanism could be the fact that these infections are chronic and therefore promote the development of an oligoclonal TCR repertoire characterized by preferential use of specific V and J gene segments (Attaf *et al*., 2018; Jamaleddine *et al*., 2024). For example, public TCRs, shared across multiple individuals, are shown to be a hallmark of CMV-specific immunity (Attaf *et al*., 2018; Schober *et al*., 2020). Some individuals develop "superdominant" or "supra-public" TCR clonotypes, which have a high probability of generation and can represent a substantial proportion of all CD8+ T cells (Attaf *et al*., 2018; Schober *et al*., 2020). In this context, the amplification of public TCR clones (Emerson *et al*., 2017) would blur the HLA-based signal and therefore add noise in the context of our predictive task. A potential future direction to address this would be to test whether removing extremely high-frequency TCRs from the analyses improves model performance; however, we do not explore this possibility in the current manuscript.

A key finding of our study is the strong association between *HLA-TCR coherence* and aGVHD severity. Higher HLA-TCR coherence, defined as a strong negative correlation between TCR repertoire feature dissimilarity and sharing more HLA alleles, was linked to more severe aGVHD outcomes. This suggests that individuals with highly coherent TCR repertoires may be predisposed to stronger alloreactive immune responses post-transplant. One possible mechanistic interpretation is that in individuals with high HLA-TCR coherence, TCR selection during thymic development is more tightly shaped by the host’s HLA environment, resulting in a repertoire finely tuned to recognize slight variations in HLA-presented peptides. Post transplant, when T cells encounter mismatched or semi-mismatched recipient HLAs, this heightened sensitivity could amplify alloreactivity, thus increasing GVHD risk. Notably, a predictive model, which incorporated HLA-TCR coherence alongside classical HLA-based metrics such as peptide binding patterns (Reynisson *et al*., 2020) and HLA evolutionary distances (Di *et al*., 2020), demonstrated that coherence was the most influential factor in predicting aGVHD severity in our cohort. This hints that TCR repertoire characteristics could offer supplementary insight but would require further evaluation in additional cohorts. This echoes the hypothesis that some individuals are simply "dangerous donors" (Baron *et al*., 2007). This idea stems from an observation that in the context of HLA haploincompatibility, not all grafts elicit aGVHD, which only highlights the unpredictable nature of aGVHD. Indeed, it has previously been observed in mice that it is possible to select for "good" or "bad" donors, through generations of crossbreeding high and low alloresponders (Cao *et al*., 2003). Our HLA-TCR coherence results suggest the relevance of potential alloreactive responses for aGVHD risks in humans, warranting further investigation into this phenomenon.

Beyond HLA-TCR coherence, our findings indicate that a donor’s infection history significantly influences aGVHD risk. We identified several previously reported viruses, such as *Herpesvirus 6B* and *7* (Phan *et al*., 2018), as well as a few novel ones including *Human coronavirus HKU1* and *Human adenovirus D*, as potential risk factors for severe aGVHD. While Human herpesvirus 4 (EBV) and 5 (CMV) seropositivity in the donor have been associated with a higher aGVHD risk (Kołodziejczak *et al*., 2021), we did not find this association to be statistically significant in our data. This is not unexpected since the association between CMV and aGVHD is conflicting int he literature (Cantoni *et al*., 2010; Ghobadi *et al*., 2019; Giménez *et al*., 2019; Teira *et al*., 2016). In contrast, we found that a history of infection with *Human adenovirus B* and *C* as well as *Orf virus* and *Herpesvirus 3* appeared protective. This suggests that imprints of prior immune challenges imprint lasting effects on the donor’s TCR repertoire, influencing post-transplant immune responses. A molecular mechanism at the root of this observation could be that these infections amplify virus-specific clones that then make their way into the graft. The difference between the risky vs protective viruses could be a difference in alloreactivity or cross-reactivity once in the recipient (Barrett, 2011). The association between infection history in the donor and tissue manifestations of aGVHD remain exploratory and need further validation, especially given that a VirScan hit can indicate not only a prior immune response from infection or vaccination to that virus but also potential cross-reactivity with related antigens. To further dissect the relationship between infection history and TCR repertoire composition, we developed a computational framework to identify virus-associated TCRs. By leveraging classifiers trained on VirScan profiles, we successfully isolated a subset of TCRs with statistically significant viral associations. The majority of these “vaTCRs” were specific to a single virus label, while a subset exhibited cross-association. Notably, vaTCRs linked to "dangerous" viruses (i.e., viruses associated with increased aGVHD risk in our study) were enriched in donors whose recipients developed severe aGVHD. Conversely, vaTCRs associated with "protective" viruses (i.e., viruses associated with lower aGVHD risk in our study) were negatively correlated with aGVHD severity. These results provide a potential mechanism through which prior infections shape immune outcomes in transplantation settings. The specifics of the mechanisms underlying these associations and their link to aGVHD occurrence remain to be fully elucidated.

Finally, we integrated our multi-modal data using a deep learning model stratifying the individuals by risk of developing aGVHD. Upon examining the model’s performance by iteratively masking input data modalities, we found that the TCR repertoire features (e.g., the V- and J-gene usages, and CDR3 amino acid statistics) are the single most important data modality to include. Moreover, our findings indicate that simply relying on clinical information such as sex and age mismatch is not enough to build a statistically significant risk stratification. Indeed, we find that all the types of data appear to be additive in their contribution to the risk prediction. Finally, our analyses show that the expressivity of deep learning models allows us to model multivariate data from various modalities in the context of survival analysis.

Several alternative strategies were explored during this study, though they ultimately proved ineffective for the analyses presented here. We initially attempted grouping individuals based on a binary presence/absence vector of each HLA allele rather than clustering on structural features such as clefts; this approach led to poor generalization, particularly for rare alleles making their way into the test set. Standard, non-contrastive learning methods were applied to predict TCR features from HLA or VirScan data. However, because individual repertoires are overwhelmingly similar and traditional regression loss functions such as mean squared error or absolute error are optimized to minimize absolute prediction errors, fail to capture the subtle distinctions that encode meaningful biological variation. This limitation motivated the adoption of contrastive learning methods, which are specifically designed to emphasize relative differences between similar samples rather than relying on absolute values. Direct classifiers for predicting aGVHD grade from repertoire or VirScan data also failed, as the relevant signal was temporally encoded, motivating the use of survival models and specifically multivariate CoxPH. Finally, Fisher’s exact tests for vaTCR enrichment, as previously implemented by (Emerson *et al*., 2017), were hindered by stringent multiple testing corrections (corrected p-value = p-value / # of tests) from sheer number of TCRs tested (>1M), preventing any TCRs from reaching significance in VirScan association. Collectively, these negative results informed the methodological choices made for the final analyses in this manuscript.

Overall, our findings have significant implications for the field of transplantation immunology. First, they highlight the limitations of traditional HLA matching in fully capturing the complexity of T cell alloreactivity. Incorporating TCR repertoire analysis and coherence metrics into donor selection strategies may be a useful albeit expensive consideration for refining aGVHD risk stratification. It is important to note that cheaper to obtain surrogates for overall viral exposure such as donor age were found to be bad predictors of aGVHD risk. Second, our results suggest that pre-transplant screening of donor infection history might help inform assessments of transplant compatibility. Finally, our computational approach for isolating vaTCRs offers a novel tool for studying virus-specific immune responses in various clinical contexts, including autoimmune diseases and vaccine development.

While the cohort provides a rare resource, the study was conducted in a previous therapeutic era (1990-2005). Because aGVHD prophylaxis today is largely centered on post-transplant cyclophosphamide (PTCy), the results must be confirmed in modern cohorts. Moreover, VirScan signatures represent past viral exposures or immunizations, and some observed associations were unanticipated. Some unexpected viral hits such as *Cowpox* or *Orf* or *Dengue virus* are likely attributable to cross-reactive antibody responses rather than true recent exposure in the cohort. As such, VirScan data should not be interpreted as seropositivity but rather as a viral immune signature. Nevertheless, even when the specific viral label reflects cross-reactivity, these signals still capture meaningful aspects of an individual’s immune history and can provide useful structure for detecting associations with GVHD risk. As a result, the pathways by which these prior infections might shape GVHD risk cannot be determined from the available data and merit additional investigation.

In conclusion, our study underscores the importance of considering both genetic and environmental factors in shaping TCR repertoires and their implications for transplantation outcomes. By integrating serological profiling, computational modeling, and TCR sequencing, we provide a framework for advancing precision medicine approaches in transplantation immunology. While our study provides compelling evidence for the role of HLA-TCR coherence and infection history in shaping immune responses post-HSCT, several questions remain. Future studies should aim to validate our findings in larger and more diverse cohorts using currently used aGVHD prophylaxis regimens based on PTCy, including more recipients of mismatched and haploidentical transplants. Additionally, mechanistic studies are needed to determine whether vaTCRs contribute directly to aGVHD pathogenesis or merely serve as biomarkers of immune risk. Finally, integrating single-cell TCR sequencing with functional assays to more robustly characterize infection history in individuals could provide deeper insights into the antigen specificity and functional properties of vaTCRs.

## MATERIALS AND METHODS

### Cohort and data collection

We analyzed the cohort of 401 healthy hematopoietic stem cell transplant (HSCT) donors described in (Emerson *et al*., 2017) (Table S1), all of whom donated grafts for recipients. Donor demographics, including age, sex, ethnicity, race and infection history, were obtained, as well as recipient preparative regimen and aGVHD prophylaxis. T cell receptor (TCR) repertoire sequencing data was downloaded from Adaptive Biotechnologies website linked in the original manuscript (Emerson *et al*., 2017). Infection history information obtained from VirScan assays for donors was downloaded from (Manuscript in preparations for submission to *Datasets*). Details on VirScan procedure can be read in (Bender Ignacio *et al*., 2019; Walti *et al*., 2021), and data were analyzed with *phippery* (Galloway *et al*., 2023) using the z-score methodology described by (Mina *et al*., 2019), with a threshold for positive detection of a response to each peptide (hit) set to a z-score ≥ 7.0 for both of two replicates. Because of the overlapping peptide design of the VirScan library, multiple hits within a virus that overlapped by 7 or more amino acids were flattened into a single hit, preserving the sequence with the highest average z-score across replicates.

### HLA clustering and binning

HLA alleles were grouped based on structural similarity and peptide-binding preferences in accordance with the established hierarchical clustering methods (Pierini and Lenz, 2018; Thomsen *et al*., 2013). Specifically, for each of the six allele categories (A, B, C, DRB1, DP, DQ), we performed k-means clustering over each allele’s peptide binding cleft sequence, using a Grantham distance (Grantham, 1974) between amino-acid residues (see Figure S1). For all analyses in the manuscript, HLA alleles are always grouped according to their peptide binding cleft similarity.

### TCR repertoire selection models

For each individual TCR repertoire in our cohort we trained a selection model, using the soNNia software described in (Isacchini *et al*., 2021; Sethna *et al*., 2020a). Specifically, this model characterizes how sequence features of a functional TCR (e.g., V-, J-gene usage, CDR3 length and its amino acid composition) in a repertoire are distinct from the expected statistics shaped by the generation process by VDJ recombination; these differences may be due to thymic selection or selection for antigen specificity (hence the term *selection model*). Thus, the probability P_!"#$_(σ) to observe the TCR sequence σ in a functional repertoire is described by reweighing the receptor’s pre-selection generation probability 𝑃_%&’_(σ) with sequence-specific selection factors 𝑄(σ) (Figure 1B): 𝑃_()*+_(σ) =, 𝑄(σ) × 𝑃_%&’_(σ), where 𝑍 is the appropriate normalization factor. For each individual, the generation model 𝑃_%&’_ is learned from the unproductive repertoires, using the IGoR software (Marcou *et al*., 2018), and the selection factors are learned by training on an individual’s functional repertoire, using a maximum likelihood approach from the linear soNNia model (Isacchini *et al*., 2021; Sethna *et al*., 2020a). The learned selection factors can be represented by a vector, where each entry reflects the deviation for the preference of a given sequence feature in the functional repertoire over the bias from the pre-selection generation process. Each selection vector (*Q*) consists of 1797 entries for possible V, D, and J genes, CDR3 lengths, and amino acid compositions across the CDR3 region, and the selection contribution 𝑄(𝜎) for a given sequence 𝜎 is the product of *Q*-factors associated with the specific features of that sequence; see (Isacchini *et al*., 2021; Sethna *et al*., 2020a) for more details.

We train generation and selection models separately for each individual in the cohort, and use them to evaluate the probability of specific sequences σ from a given individual, and to characterize individual-specific selection factors (*Q* factors, Figure 1B); we compare selection models across individuals to characterize the pairwise distances between their TCR repertoires.

### Predicting differential TCR repertoire compositions from HLA and VirScan features, with contrastive learning

We aim to assess how differences in the HLA composition and/or infection history between two individuals determine the differences between the composition of their TCR repertoires. To do so, we first represent an individual 𝑖 by a feature vector 𝑥. ∈ ℝ^/^ denoting their HLA haplotype information and/or infection history from VirScan assay readout; this vector has dimension 𝐷, corresponding to the input dataset dimensionality.

Next, we represent the difference between two TCR repertoires 𝑖 and *j* by the differences in their corresponding selection models (i.e., by the difference in the logarithm of their feature-specific selection factors (𝑄. and 𝑄_0_) as: Δ log 𝑄_.0_ = log 𝑄. − log 𝑄_0_, where Δ log 𝑄_.0_ ∈ ℝ^,121^.

To assess the impact of HLA haplotype and/or infection history shape the TCR repertoire, we train a contrastive neural network with a single encoder (Bromley *et al*., 1993) that takes a pair of individuals’ HLA/VirScan feature vectors (x., x_0_) as input, and predicts the difference in their repertoire’s selection factors, Δ log 𝑄_.0_. A single encoder 𝑓_3_ is applied to each person’s HLA/VirScan feature vector, producing latent representations 𝑓_3_(𝐱.) and 𝑓_3_>𝐱_0_?; their difference 𝐲A_.0_ = 𝑓_3_(𝐱.) − 𝑓_3_>𝐱_0_? is then regressed against the empirical Δ log 𝑄_.0_. We optimize the model by minimizing the mean squared error (MSE) between 𝐲A_.0_ and Δ log 𝑄_.0_, thereby forcing the encoder to place individuals with similar selection factors close together in latent space, while pushing dissimilar ones farther apart.

We trained the network on *N* = 10,000 samples of randomly paired individuals, with a loss function,

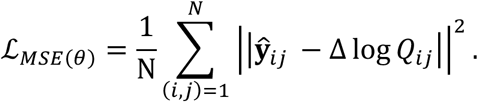

The encoder network 𝑓_>_ is a fully connected feedforward neural network with the following architecture details:

- Linear transformation : ℝ^/^ → ℝ^?,:^
- Activation ReLU (Rectified Linear Unit)
- Output layer: Linear transformation from: ℝ^?,:^ → ℝ^,121^

This design encourages the encoder to map input samples into a learned feature space where meaningful differences between pairs can be captured by simple vector subtraction.

Optimization was performed using the Adam optimizer with a learning rate of 10^@A^and a batch size of 64. The model was trained for 100 epochs. Model was implemented in PyTorch (v2.5.1) and executed on a MacBook Pro M1 laptop.

### Model evaluation

To avoid overfitting, we held out 20% of individuals in our cohort as an external test set (none of these individuals contributed to the 10,000 training pairs). To evaluate our model’s performance and generalization, predictions using the trained encoder were computed on pairs formed from this held-out test set. The training set consisted of 10,000 pairs of individuals selected from the training individual pool. A 20% validation set used for hyperparameter tuning was selected at random from the 10,000 pairs. In other words, the 10,000 pairs were split 8000:2000 between training and validation sets. The test set used for evaluation of the model’s performance was 10,000 pairs of individuals formed between a test set individual and a training set individual. This setup for the test set mimics the situation where for given a pre-trained model, we obtain a new individual and encode their features into our pre-existing pre-trained encoder map. For all instances of the contrastive neural network, performance was measured via Pearson correlation coefficient between the predicted and the actual values for the repertoire features.

### TCR presence/absence prediction

We constructed a dataset comprising 𝑁 = 10,000 paired samples, where each pair (*i, j*) is represented by two input feature vectors of HLA cleft clusters and/or Virscan assay readouts >𝐱., 𝐱_0_?. For each individual *i*, we derived a binary target vector 𝐭. ∈ {0,1}^B^, where each element indicates the presence (1) or absence (0) of a specific TCR among a predefined set of 𝐾 = 1000 unique TCRs. These TCRs were selected at random from a pool of private, public or superpublic TCRs. Private TCRs are those seen in less than 50 individuals in our cohort, public TCRs are seen in 50-300 individuals, and superpublic were defined as seen in at least half of our cohort (>300). For each pair of individuals, we calculated the vector Δ𝐭_.0_ = 𝐭. − 𝐭_0_, defined as the difference in presence or absence of the random set of TCRs.

Next, we trained the previously described contrastive neural network to predict the difference between TCR presence /absence Δ𝐭_.0_ of a pair of individuals from the inputs. The training and evaluation schemes are otherwise similar to our modeling of differential TCR repertoire compositions from HLA and/or VirScan features.

### Model feature importance

To evaluate the relative importance of each input feature, we partitioned the various input features into groups (M) by input type: (HLA cleft clusters, HLA types, Virus groups, Clinical data, etc.). For each model training run, we selected a subset of these feature groups by masking them (placing their values to zero in the inputs). The binary vector 𝑚 ∈ {0,1}^4^ indicating which masked (0) or unmasked (1) inputs (M) were used as well as the performance of that specific training run was recorded 𝑢 ∈ ℝ. We repeated this procedure until all possible combinations of input groups were explored, including single input groups. To quantify the relative contribution of each group to the overall performance, we trained a linear regression model using the binary vectors 𝑚 ∈ {0,1}^4^ as inputs and the run performances 𝑢 ∈ ℝ as outputs.

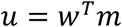

where 𝑤^C^ ∈ ℝ^4^, corresponds to the learned weights (coefficients) of the linear regression and the relative importance of each feature to the performance of the model. A large absolute value of the coefficient indicates a high importance and a value close to zero (positive or negative) indicates a lesser importance.

### SHAP factors for model interpretability

To interpret our models, we applied SHapley Additive exPlanations (SHAP), an approach from the field of game theory that estimates the marginal impact of each player on the game outcome given all the other players involved (Lundberg and Lee, 2017). In our context, we use it to calculate each input feature contribution on the model output. SHAP values were computed using a kernel-based approximation method adapted for deep learning models. We used the python package SHAP (v0.47.2) for these calculations (Lundberg and Lee, 2017).

### Selection-specific Jensen-Shannon divergence between TCR repertoires

To characterize how differential selection shapes differences between repertoires, we computed the Jensen-Shannon divergence 𝐷 >𝑃^E^ , 𝑃^E!^for all repertoire pairs (𝑟, 𝑟^F^), each characterized by their post-selection models 𝑃^E^ and 𝑃^E!^ , inferred by the soNNia software (Sethna *et al*., 2020b).

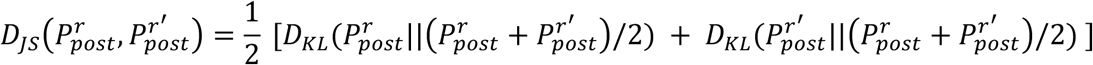

where 𝐷_BG_(𝑃, ||𝑃: ) = ⟨log: ^H"(I)^ ⟩ is the Kullback-Leibler distance between two distributions.

As noted previously, for a given repertoire 𝑟, we model the probability 𝑃^E^ (𝜎) to observe the TCR sequence σ in a functional (post-selection) repertoire by 𝑃^E^where 𝑃^E^ (𝜎) is the generation probability reflecting the recombination process, and 𝑄^E^(𝜎) is the sequence-specific selection factor; for brevity, the overall normalization is absorbed into 𝑄^E^.

Accordingly, differences between two repertoires (𝑟, 𝑟^F^) can be decomposed into (i) generative differences–captured by differential 𝑃_%&’_’s (e.g., due to distinct V/D/J gene or allele usages, insertion/deletion statistics)– and (ii) selection differences–captured by differential 𝑄 factors (e.g., due to distinct HLA compositions, antigenic exposures, or other immunological pressures.)

To factor out the effect of generative process, we first infer a separate generation model 𝑃^E^ (𝜎) for each repertoire 𝑟. We then construct the four “hybrid” post-selection distributions 𝑃^(.,0)^(𝜎) = 𝑃. (𝜎)𝑄^0^(𝜎), with 𝑖, 𝑗 ∈ {𝑟, 𝑟^F^}, i.e., we pair each generation model with each selection model, and define 𝑟^(.,0)^ as the sequences generated by the hybrid model 𝑃^(.,0)^.

We then define a modified selection-specific Jensen-Shannon divergence that averages the divergence between the two selection models while holding the generation model fixed, and then averages over the two possible generation models,

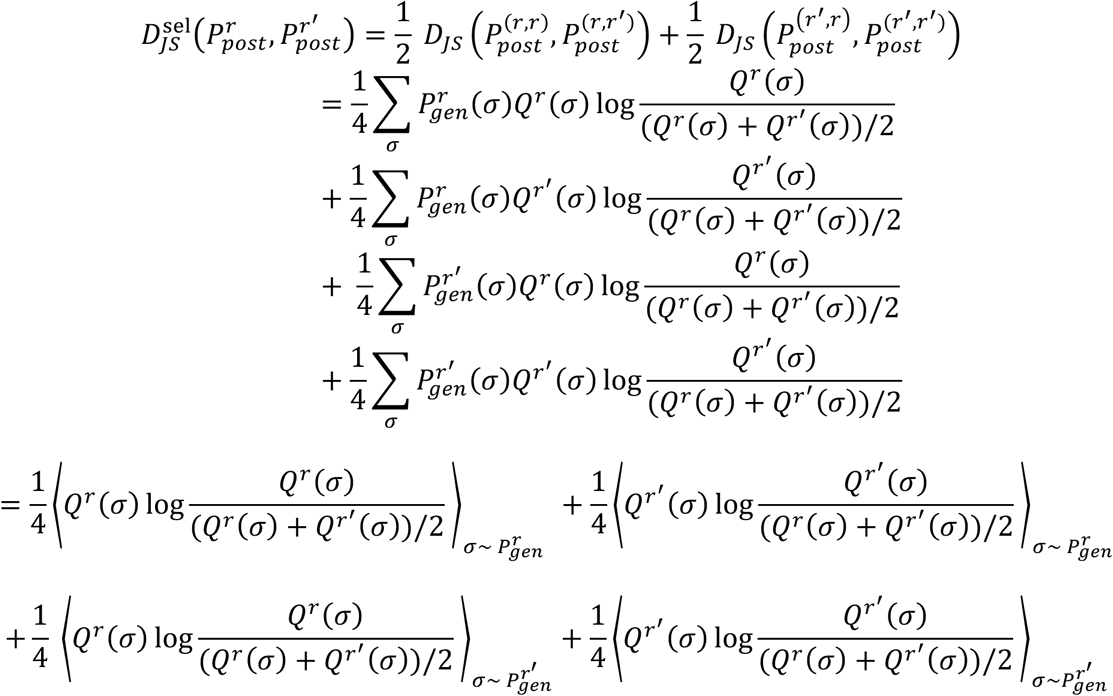

where the ⟨. 〉 ( denotes the expectation of the argument over sequences sampled from the generation model 𝑃, specified in the subscript.

The resulting selection-specific Jensen-Shannon divergence is computed for all repertoire pairs, quantifying the extent by which differential selection differences–independently of generative differences–drive compositional divergence between repertoires.

### HLA-TCR Repertoire Coherence

To quantify HLA-TCR repertoire coherence, we calculated the correlation between DJS values and the number of shared HLA alleles across donor pairs. Specifically, an individual’s HLA-TCR coherence is the Pearson correlation coefficient between DJS values and numbers of HLA alleles in common. We used the p-value of the Pearson correlation as a criterion to select only the HLA-TCR coherences that were statistically significant. Next, binning the donors into quartiles by the HLA-TCR coherence, we classify individuals in Q1 as "coherent" individuals. Similarly, individuals from Q4 were classified as the least coherent.

### Random Forest model for graft outcome prediction from HLA features

To evaluate the predictive power of HLA-TCR repertoire coherence, we trained a Random Forest regressor. As inputs we used HLA-TCR coherence, number of predicted binding peptides as well as allele evolutionary distances, as detailed below.

To characterize each individual’s antigen presentation capacity, we ran NetMHCpan-4.1 (class I), the widely validated neural network predictor for pan-allelic HLA presentation (Reynisson *et al*., 2020). We downloaded complete proteomes for ten selected viruses (Supplementary Table II) from the UniProt database (UniProt Consortium, 2025) and generated a database of all possible 8-12-mer peptide from all proteins. NetMHCpan scored each peptide against every class-I allele observed in our cohort; peptides ranked within the top 0.5% for an allele were designated as strong binders(Reynisson *et al*., 2020). Summing these predicted binders across the two present alleles at each HLA-A, -B, and -C locus produced, for every individual, these numbers capture the total repertoire of high-affinity viral peptides that their class-I HLA alleles are predicted to present (Figure S2A-B ).

To characterize the intra-HLA allele diversity for each individual, we calculated the divergence between each pair of class I HLA alleles for each individual in two distinct ways. (i) *Evolutionary distance* was taken as the pairwise amino acid divergence of the peptide binding clefts for every HLA-A, -B, and -C allele pairs, following the approach described in (Chowell *et al*., 2019; Pierini and Lenz, 2018); see Figure S2C-E. (ii) *Functional distance,* in contrast, compares the predicted strong-binding peptide repertoire to be presented by each allele. A functional distance of zero means there is no difference between the bound peptides for both alleles, pointing either to a homozygous individual for that locus or two alleles that bind the exact same peptide pool. This measure differs from the evolutionary distance by comparing peptide binding rather than directly the binding cleft. Indeed, two different clefts could end up binding the exact same pool of peptides (Hertz and Yanover, 2007; Sidney *et al*., 2008).

Next, the HLA-TCR coherence, the number of HLA’s in common, as well as the total numbers of predicted peptides bound, functional and evolutionary distances for each of HLA-A, HLA-B and HLA-C were used as inputs to predict the graft outcome using a Random Forest Regressor. The graft outcome is a number between 0 and 4, that corresponds to the grade (severity) of acute graft versus host disease (aGVHD) developed by the patient after the HSCT procedure (grade 0 meaning no aGVHD, grade 1-2 meaning mild aGVHD and grades 3-4 meaning severe aGVHD) (Glucksberg *et al*., 10 1974).

To evaluate the random forest model, we held out 20% of individuals as a test set and performed this procedure 10 times. For feature importance evaluation, we calculate its contribution to reducing the gini impurity within the decision-making process (Louppe *et al*., 2013) as well as a permutation importance test (Altmann *et al*., 2010).

### Survival analysis for VirScan and aGVHD

Cox proportional hazards (Cox PH) modeling was used to train a combined model for all viruses in VirScan and their association to aGVHD occurrence. Only donors with both VirScan and clinical information were included in the training procedure. Multivariate models were used to correct for known predictors of aGVHD (donor and recipient sex mismatch, donor age, CMV serostatus pairing, aGVHD conditioning and prophylaxis regimens) and directly compare them to the VirScan data predictive potential. Univariate models were also assessed on single viruses as well as Kaplan-Meier survival curves with log-rank test (Figures S4-S5). Multivariate CoxPH model was considered statistically significant when overall p-value was <0.05 and specific virus associations to aGVHD occurrence were considered when corrected p-value was below 0.05. We categorized viruses as "dangerous" or "safe" based on their hazard ratios and confidence intervals and direction of association from CoxPH model results.

### Identification of Virus-Associated TCRs (vaTCRs)

We developed a computational pipeline to identify virus-associated TCRs (vaTCRs) from the donor repertoires. TCRs were filtered through three sequential steps:

1. **Shared TCR Selection:** We retained TCRs found in at least two individuals, reducing the dataset from 80 million to ∼2 million unique sequences.
2. **Predictive Association Filtering:** Logistic regression models were trained to predict TCR presence based on VirScan infection profiles, where VirScan profile vectors served as input and the presence or absence of a TCR was the output for each logistic regression model. TCRs with significant predictive performance on validation sets (p-value < 0.05, AUC-ROC > 0.6) were retained (∼10,000 sequences).
3. **Virus-Specific Enrichment Analysis:** To associate each retained TCR with a specific virus, contingency tables were constructed based on the VirScan positivity threshold (>=3) and the presence or absence of each TCR. Fisher exact tests were then applied, and following multiple-testing correction significantly enriched virus-TCR pairs were identified (∼4,577 vaTCRs).

In order to calculate a false discovery rate (FDR) on our entire computational pipeline, we shuffled the VirScan infection profiles between the donors, before running steps 2 and 3. On Step 2, for a *true hit* we calculate the difference in mean AUC-ROC between the predictor trained on the true VirScan vectors and the predictor trained on shuffled VirScan vectors. For a *false hit*, we calculate the difference in mean AUC-ROC between two predictors trained on shuffled data (shuffled separately). For both instances we only retain *hits* that have an AUC-ROC of >0.6 and for which the difference in AUC-ROC is statistically significant (p-value<0.05). Then, Fisher tests from Step 3 are run on both *true* and *false hits*, to calculate an FDR. Figure S7A shows the distribution of AUC-ROC scores as well as p-values between the *true* and *false hits* for a randomly selected set of 10,000 sequences. The bottom right corner represents the filter at Step 2: AUC-ROC >0.6 and a statistically significant difference compared to a control shuffled set. Figure S7B shows the calculated number of false hits for Step 2 and Step2+Step3 as the number of false hits / the number of TCR evaluated for 25 randomly selected data slices.

### TCR sequence clustering

To characterize vaTCRs, we computed pairwise distances using tcrdist3 (Mayer-Blackwell *et al*., 2021) and visualized these distance relationships using Uniform Manifold Approximation and Projection (UMAP) (Becht *et al*., 2018). K-means clustering was applied to group sequences in space and to detect sequence motifs, and cluster-specific amino acid patterns were visualized using sequence logo plots created with the *logomaker* python package (Tareen and Kinney, 2019). Next, we used *tcrpheno*, to classify each sequence into the four distinct T cell states as described in (Lagattuta *et al*., 2024). For each cluster, we obtained the proportion of the various T cell state classifications.

### vaTCRs sharing in the cohort and generation probabilities

To calculate vaTCR sharing, for each sequence we compute the proportion of individuals in the *Emerson* cohort with the exact matching vaTCR amino-acid sequence in their repertoire (regardless of the annotated V gene). For each vaTCR, generation probabilities were calculated with the *OLGA* software (Sethna *et al*., 2019).

### Deep transformer for Survival Modeling

We implemented a Transformer-based deep learning model in PyTorch to predict survival outcomes. The architecture draws inspiration from the existing model *DeepSurv* (Katzman *et al*., 2016). Specifically, the input feature vector for each individual was first projected into a 64-dimensional latent space via a linear embedding layer. This embedding was then processed through a two-layer Transformer encoder with four attention heads per layer. The Transformer enabled the model to learn contextual relationships among input features through self-attention mechanisms, which is important for the various HLA inputs. The output of the Transformer was pooled and passed through a final linear layer to produce a single scalar value: *the risk score*. This *risk score* is a continuous scalar output from the final layer of the model, representing the log-risk of developing aGVHD for each patient. A higher risk score indicates a higher hazard and therefore a greater likelihood of experiencing the event (aGVHD) sooner. The risk scores do not represent probabilities directly, but rather a relative ranking of patients’ hazard under the Cox proportional hazards assumption (Harrell, 2001). These scores were used both for optimization of the Cox partial likelihood and for downstream stratification of patients into high- and low-risk groups. The model was trained using the negative Cox partial likelihood loss ℒ_𝒞ℴ𝓍_, which is suitable for right-censored data (Katzman *et al*., 2016). After sorting individuals in descending order of survival time, the loss is defined as

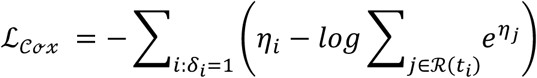

where 𝜂.is the predicted risk score for individual 𝑖, 𝛿.∈ {0,1} is the event indicator (1 for event observed, 0 for censored), and ℛ(𝑡.) is the risk set of individuals still under observation at time 𝑡.. The indices 𝑖 run over the individuals who experienced the event, 𝛿.= 1. The indices 𝑗 run over the individuals still at risk at the time 𝑡_!_.

We iteratively masked all inputs, generating all the possible combinations of inputs from the GVHD risk graph (Figure 4B). For each set of inputs, we trained an iteration of the deep learning survival transformer model using the Adam optimizer with a learning rate of 0.001 for 50 epochs on a randomly selected 80% of the samples in our cohort. To evaluate model performance on the held out 20% of samples, we computed the concordance index (C-index), which measures the proportion of all patient pairs in which the predictions and outcomes are concordant:

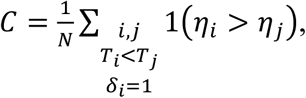

Where 𝑁 is the number of pairs, 𝑖 and 𝑗 are the compared individual indices, 𝑇.is the observed survival time for subject 𝑖. Specifically, for each comparison, we consider pairs (𝑖, 𝑗) where subject 𝑖 failed before 𝑗 and 𝑖 ‘s failure was observed. Here, a correct prediction would be a predicted score 𝜂. greater than 𝜂_0_, in other words, the model predicts a higher risk for individual 𝑖 who survived less than 𝑗. Overall, the C-index represents the fraction of such correct predictions. Thus, a C-index of 0.5 indicates random performance, while a score closer to 1.0 indicates better discriminatory power.

To visualize each model’s predictions based on the combined input modalities provided, samples were divided into high-risk and low-risk groups based on the median predicted risk score. Kaplan–Meier survival curves were generated for each group, and statistical significance between the survival distributions was assessed using the log-rank test. A significant difference between curves provides evidence that the model has learned a meaningful risk stratification and a C-index above 0.5 indicates a discriminatory power above random.

### Statistical Analysis

All statistical analyses were conducted in R and Python. Differences in model performances were tested using permutation testing and Wilcoxon signed-rank tests from the python *scipy.stats* library. Associations between infection history, TCR repertoire features, and aGVHD severity were assessed using Cox proportional hazards models (R *survival* package) and mutual information tests (python code on github). Multiple testing corrections were applied where necessary to control the false discovery rate (Yoav Benjamini and Yosef Hochberg, 1995).

## SUPPLEMENTARY MATERIALS

Figure S1 - HLA clustermaps

Table S1 – Cohort description

Table S2 – HLA cluster membership for each allele

Table S3 – VirScan viruses grouped for feature importance assessment

Figure S2 - HLA-based Feature engineering for Random Forest models

Figure S3 - Relationship between age and number of VirScan hits

Figure S4 – KM plots and univariate CoxPH for 6 viruses associated with aGVHD

Figure S5 - KM plots and univariate CoxPH for 6 randomly selected viruses

Figure S6 - vaTCR pipeline supplementary data

Figure S7 - vaTCR pipeline FDR calculations

Figure S8 - overlaps between vaTCR clusters and virus-antigen databases

Figure S9 - Feature importance from survival model saliency maps

## ACKNOWLEDGEMENTS

The authors wish to acknowledge Dr. Phil Bradley for his central role in guiding the scientific direction of this study and for ensuring its rigor. The authors acknowledge Fred Hutch Scientific Computing, supported by the National Institutes of Health (NIH) award S10OD028685 and the Fred Hutch Genomics & Bioinformatics Shared Resource, RRID:SCR_022606, supported by NIH P30 CA015704, with particular thanks to Ryan Basom for bioinformatic assistance. This work was supported by the NIH through awards R01 AI146028 (F. A. M.), R35 GM142795 (A. N.), and the CAREER award from the National Science Foundation grant 2045054 (A. N.). Additional support for A. T. was provided by the Mahan Fellowship from the Fred Hutchinson Cancer Center. This work also benefited from discussions during the 2024 program “Interactions and Co-evolution between Viruses and Immune Systems” at the Kavli Institute for theoretical physics (KITP), which is supported by National Science Foundation grant PHY-2309135, and the Gordon and Betty Moore Foundation Grant No. 2919.02. Dr. Matsen is an Investigator of the Howard Hughes Medical Institute (HHMI). This article is subject to HHMI’s Open Access to Publications policy. HHMI lab heads have previously granted a nonexclusive CC BY 4.0 license to the public and a sublicensable license to HHMI in their research articles. Pursuant to those licenses, the author-accepted manuscript of this article can be made freely available under a CC BY 4.0 license immediately upon publication.

## AUTHOR CONTRIBUTIONS

A.T., A.N. and F.A.M. conceptualized the study. A.T. performed the main analyses and result interpretation. D.H., R.B.I., T.S.A., D.Z., M.J.B., performed VirScan experiments. Z.M. and M.R. contributed to bioinformatic analyses. A.T., Z.M., A.N., and F.A.M. contributed to the analysis and interpretation of data and results. A.T. wrote the original draft and all authors reviewed, edited and approved the final manuscript.

Conceptualization: AT, AN, PB, FAM

VirScan Methodology: D.H., R.B.I., T.S.A., D.Z., M.J.B. Investigation: AT

Visualization: AT, ZM

Funding acquisition: AN, PB, FAM, AT Project administration: AT, AN Supervision: AN, PB, FAM

Writing – original draft: AT

Writing – review & editing: AT, AN, FAM, PB, ZM, MR, DH, RBI, TSA, DZ, MJB

## DATA AND MATERIALS AVAILABILITY

All code is available in the github repository : https://github.com/TrofimovAssya/TCRrepertoireforGVHD_publication. All data are available in the main text or the supplementary materials.

## SUPPLEMENTARY FIGURE LEGENDS

Fig. S1. HLA cleft clustermaps. (A-F) For each allele group, we obtained the allele binding clefts from (Reynisson et al., 2020) and calculated a pairwise distance between the clefts using a amino-acid replacement table (Grantham, 1974). Next, we performed hierarchical ward clustering and cut the hierarchy into 6 clusters for each allele group. Allele group sizes are shows directly on each plot and specific allele cluster membership is reported in Table S2.

Table S1: Cohort description

Figure S2. HLA-based feature engineering for Random Forest models. (A-B) For each MHC Class I allele in the cohort, we use (Reynisson et al., 2020) to predict binding of all possible 9-12 mers across viral proteomes from Table S4. Pannels A and B show the distributions of the sizes of sets of predicted strong binder peptides for each allele groups. (C-E) For each allele group, we obtained the allele binding clefts from (Reynisson et al., 2020) and calculated a pairwise distance between the clefts using an amino-acid replacement table (Grantham, 1974). This distance is termed HLA Evolutionary distance (HED). The functional allele distance is calculated by comparing the predicted peptide strong binders set from (A-B) using a Jaccard index. Pannels C-E show the relationship between HED and functional allele distance for each individual in the cohort. F) For each individual the HED for all three alleles is shown as a line. Individuals are grouped by pattern into clusters for better visibility using a k-means clustering over the principal components analysis of the HEDs.

Table. S2. Cluster membership for each allele Table. S3. VirScan viruses grouped by type

Table. S4. List of viral proteomes used for peptide binding predictions.

Figure S3. Relationship between age and number of VirScan hits. (A-B) For each individual in the group, A) the number of positive VirScan hits (threshold of >3) or B) the number of positive VirScan hits for aGVHD-associated viruses is compared to the individual age. Correlation and correlation significance between the two variables is assessed using Pearson correlation coefficient.

Figure S4 KM plots and univariate CoxPH models for viruses associated to aGVHD risk. Top plots: For each of the 6 viruses associated to aGVHD risk from Figure 5A, KM curves were plotted, dividing the cohort on the median VirScan value of the virus. Log-rank p-value is indicated at the top of each plot. Bottom plots: For each virus, individual VirScan values from the cohort are binned into 10 quantiles and survival times are plotted for each quantile. Top 10% and bottom 10% VirScan values are shown respectively in green and red. Univariate CoxPH hazard ratio is calculated for each virus and associated p-value is indicated on top of the plot

Figure S5 KM plots and univariate CoxPH models for control viruses. Top plots: For a random set of 6 viruses from Figure 5A, KM curves were plotted, dividing the cohort on the median VirScan value of the virus. Log-rank p-value is indicated at the top of each plot. Bottom plots: For each virus, individual VirScan values from the cohort are binned into 10 quantiles and survival times are plotted for each quantile. Top 10% and bottom 10% VirScan values are shown respectively in green and red. Univariate CoxPH hazard ratio is calculated for each virus and associated p-value is indicated on top of the plot

Figure S6 vaTCR pipeline supplementary data. (A) Barplot indicating the number of vaTCR per virus identified by the pipeline. B) Scatterplot showing the correlation between the Molluscum contagiosum and Orf virus VirScan data. 2D KDE kernel is also shown. C) For each pair of viruses sharing vaTCRs and a random set of virus pairs, the VirScan correlations are calculated and plotted as a boxplot and strip plot. Difference between correlation distributions is assessed by t-test and p-value is shown (n.s.). D) Four-way Venn diagram showing overlaps between vaTCRs associated to any coronavirus and TCR associated to any coronavirus in Mc-PAS (Tickotsky et al., 2017), VDJdb (Shugay et al., 2018) and Adaptive Biotech’s MIRA (https://doi.org/10.21417/ADPT2020COVID) databases.

Figure S7 vaTCR pipeline FDR calculations. (A) Scatterplot showing AUC-ROC values and log10 p-values for each TCR. In blue are all TCR hits from the second filter (see Methods), in orange are false hits for the second filter, in red are Fisher test-validated hits (third filter, see Methods) and in red are false hits from third filter. Numbers of TCRs in the bottom right quadrant represent the retained hits for FDR calculation (see legend). Results downscaled to show 10000 points for better visualization. B) False discovery rate (FDR) is shown as a proportion of hits / the number of TCRs tested for the second filter only (logistic regression, see Methods) and second and third filter together (logistic regression + Fisher test). To visualize FDR distribution, 25 random data slices are shown.

Figure S8 Overlaps between vaTCR clusters and virus-antigen databases. For each cluster from (Figure 6), we assigned antigen-specificity for each TCR when available, as reported by A) TCRmatch (Chronister et al., 2021) (High similarity >0.97) or B) VDJdb (Shugay et al., 2018). Proportion of virus specificity is indicated for each cluster and color-coded for each virus.

Figure S9 Feature importance from survival model saliency maps. Using the trained deep learning survival model, attention saliency maps are extracted for HLA alleles and V-J gene combinations. A) Violin plots visualize relative HLA allele importances. B) V-J gene combination feature importances are shown as a heatmap with relative importance in darker shades of blue.

