## Supplementary figures and images for "Genetic and environmental imprints on T cell receptor repertoires as predictors of graft-versus-host disease"

### Supplementary Figures

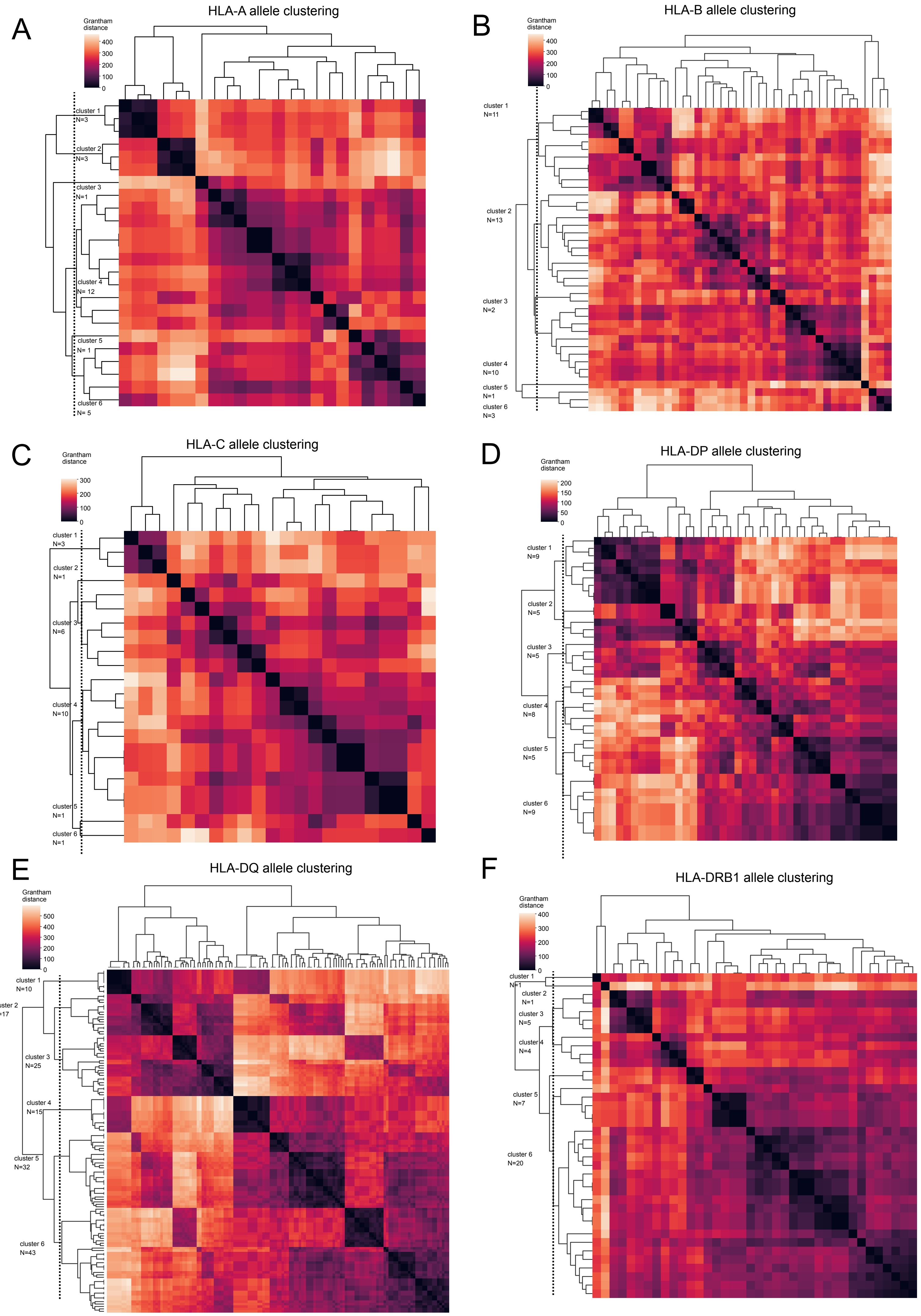

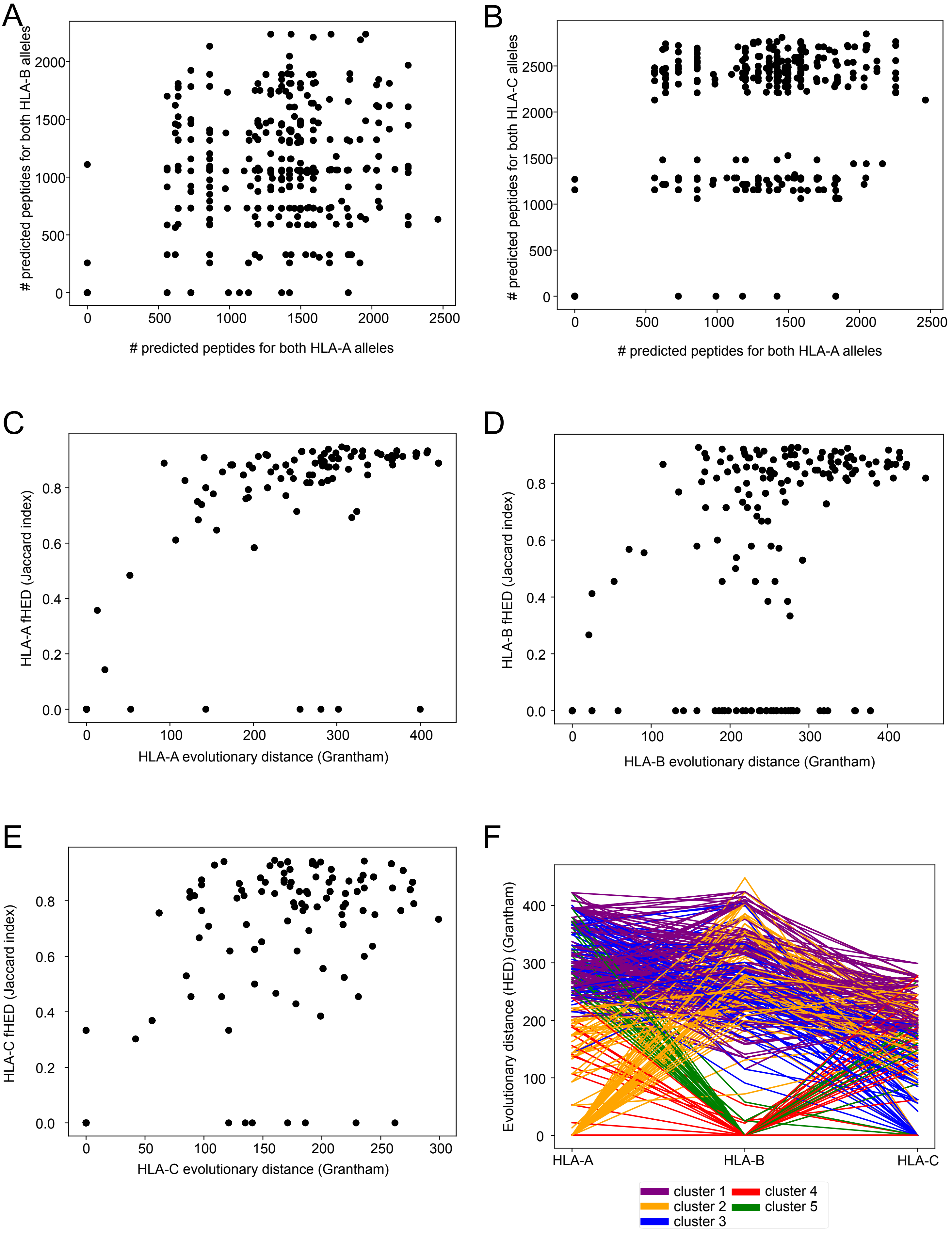

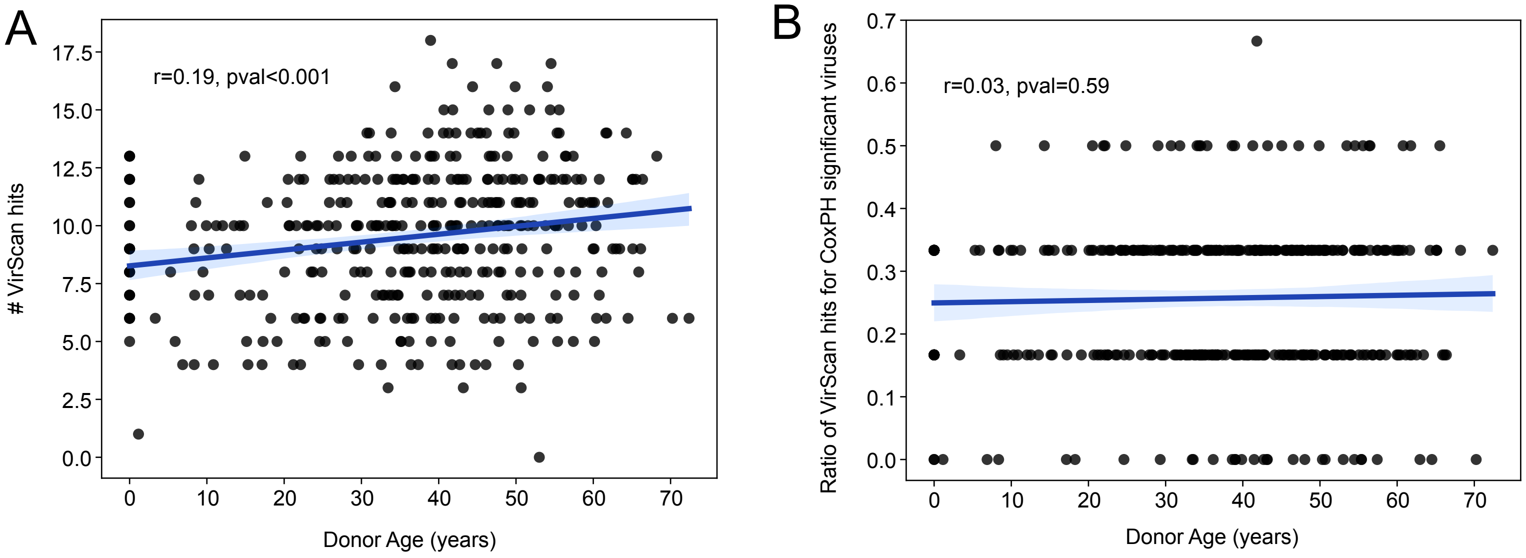

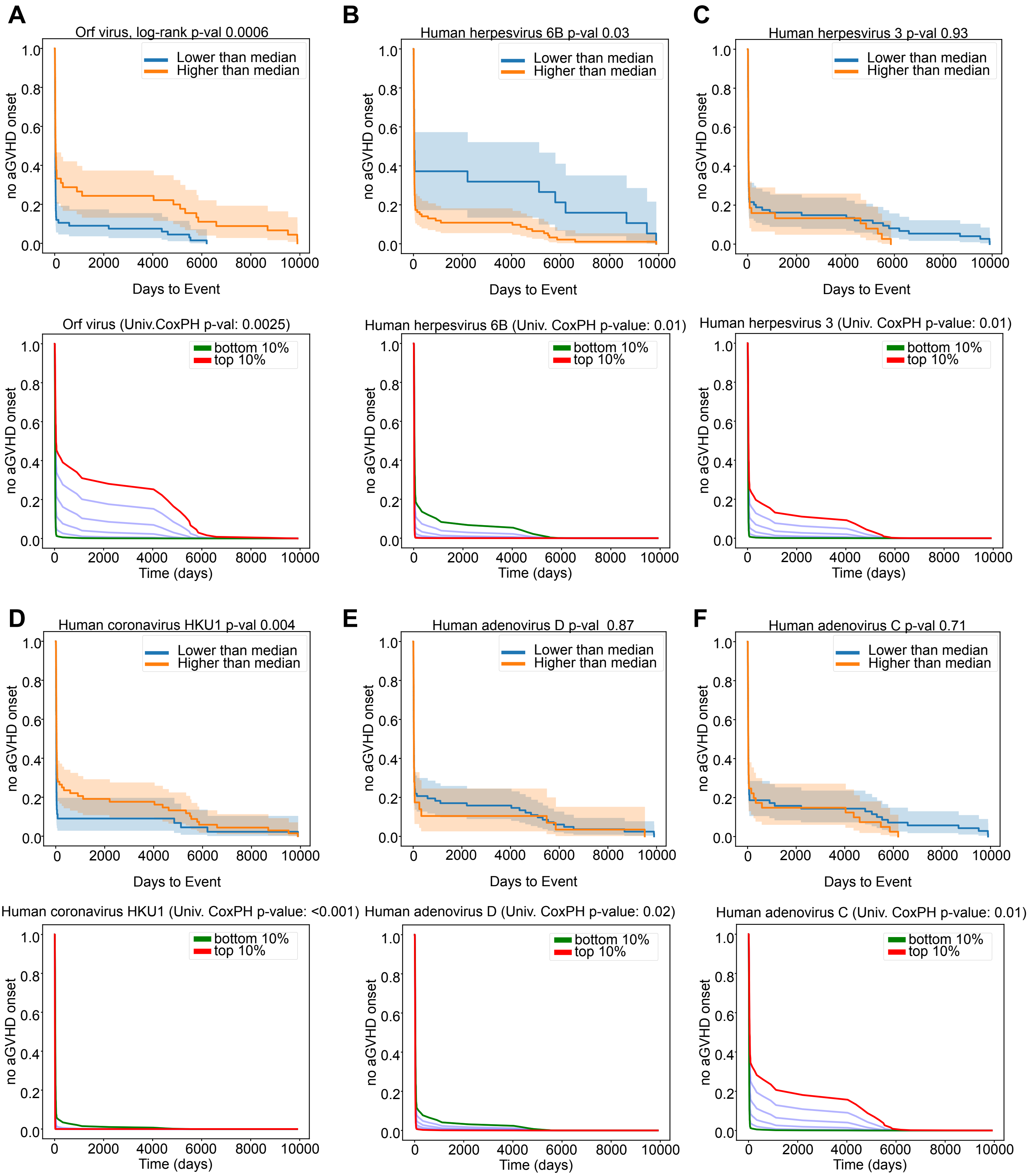

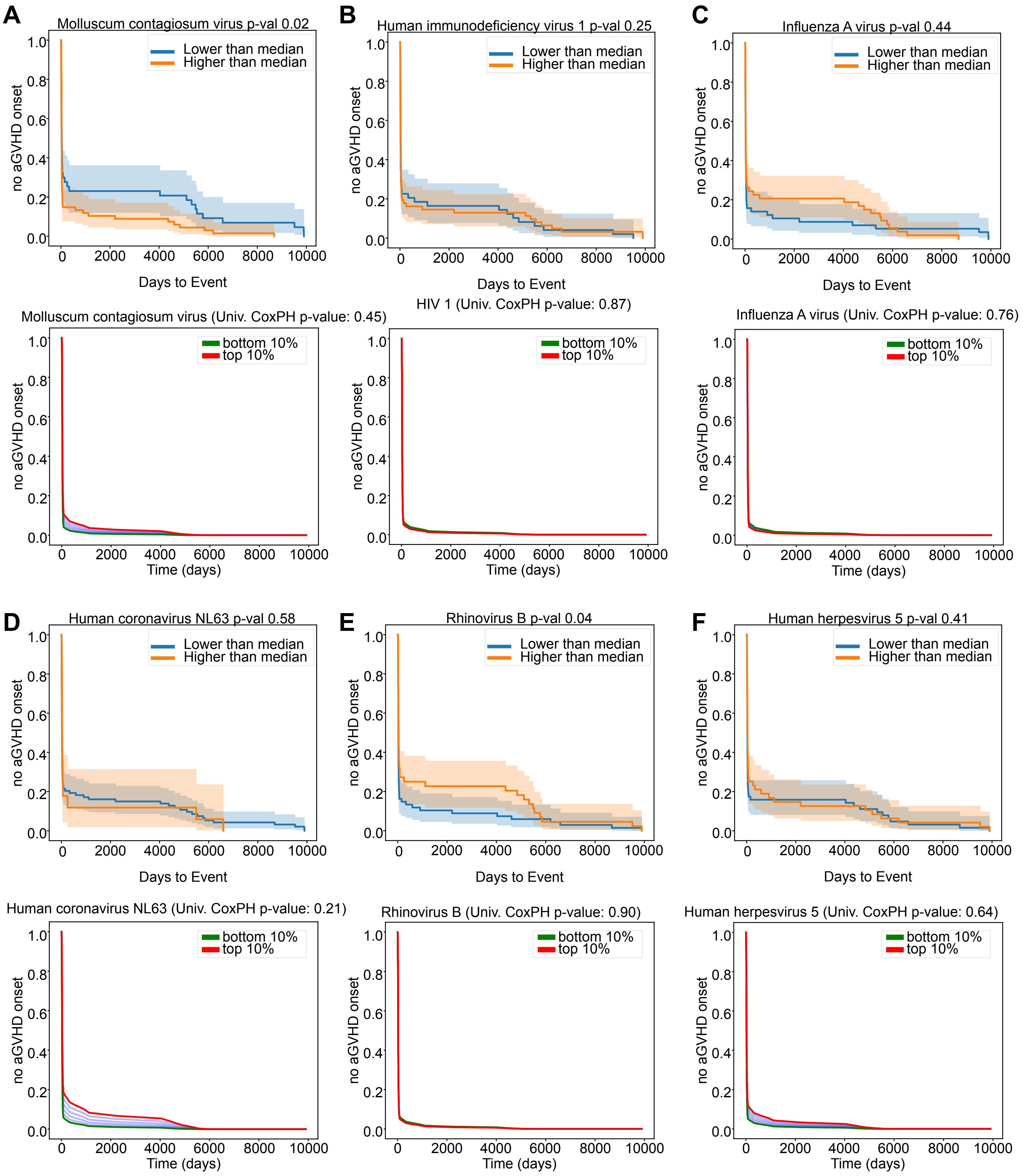

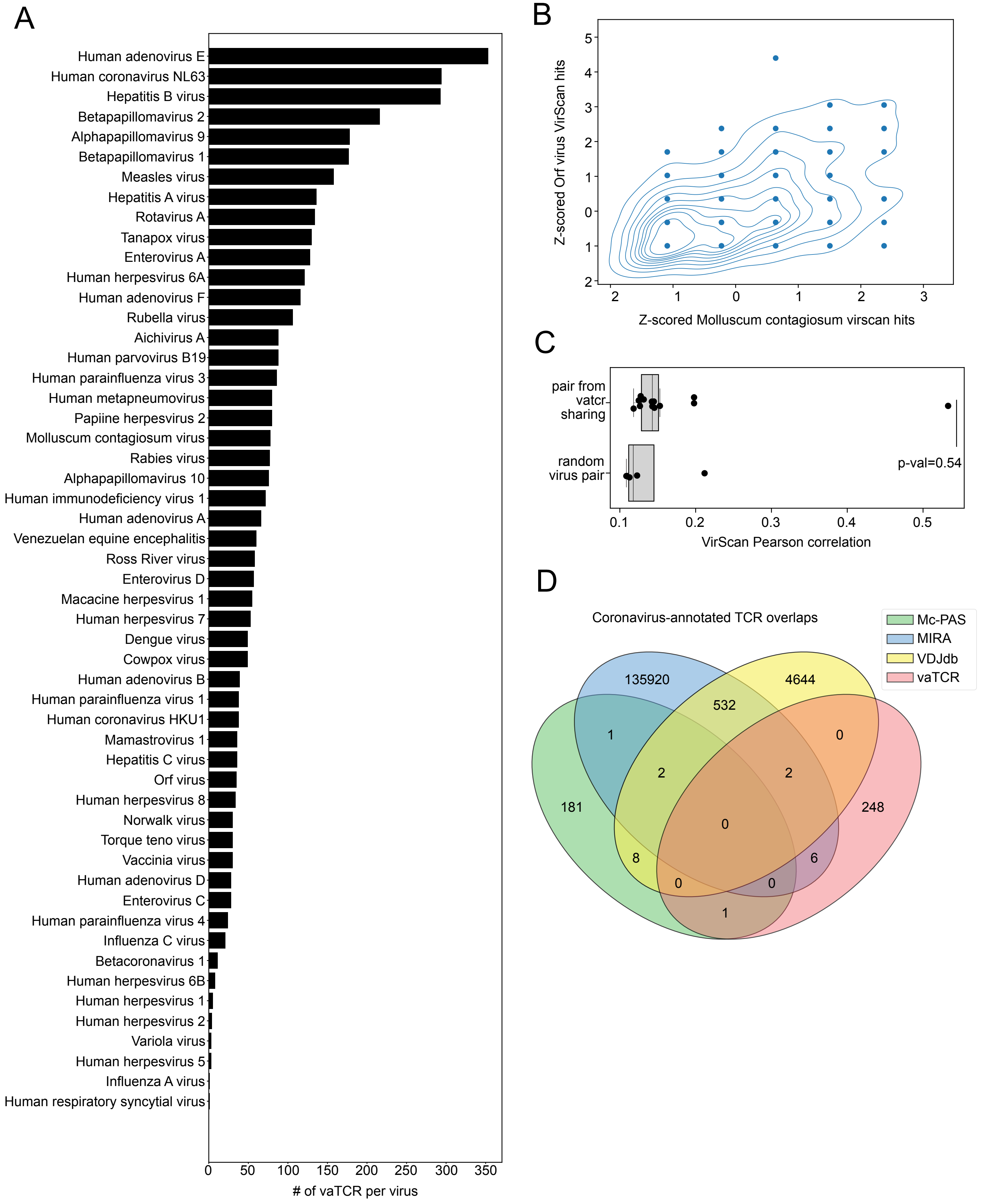

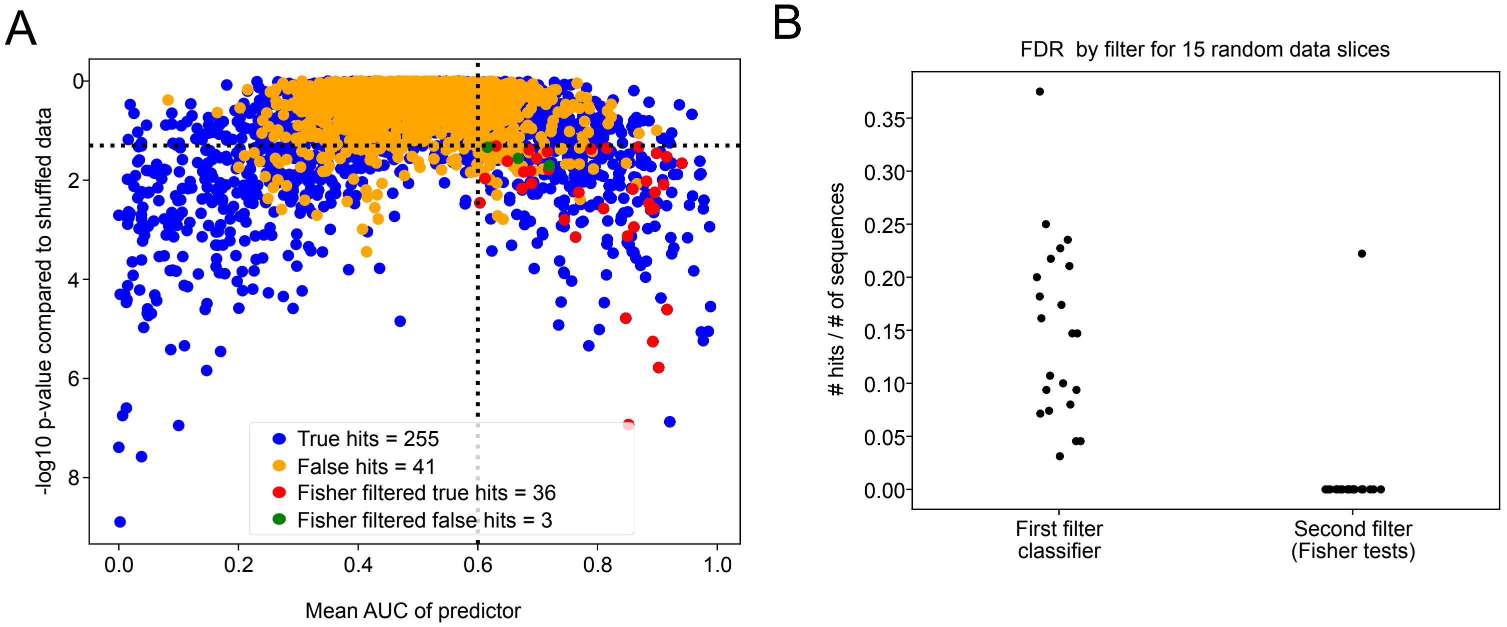

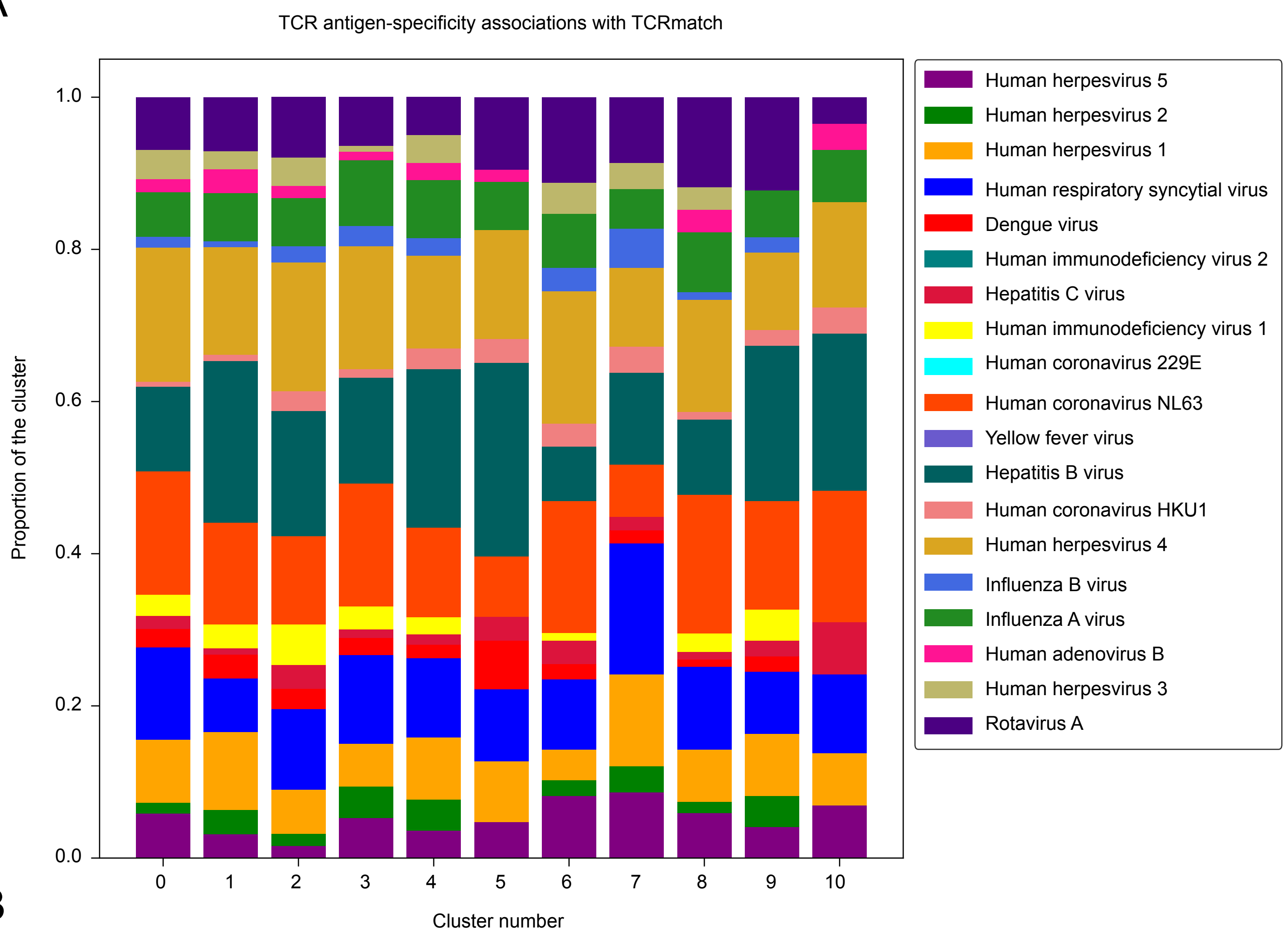

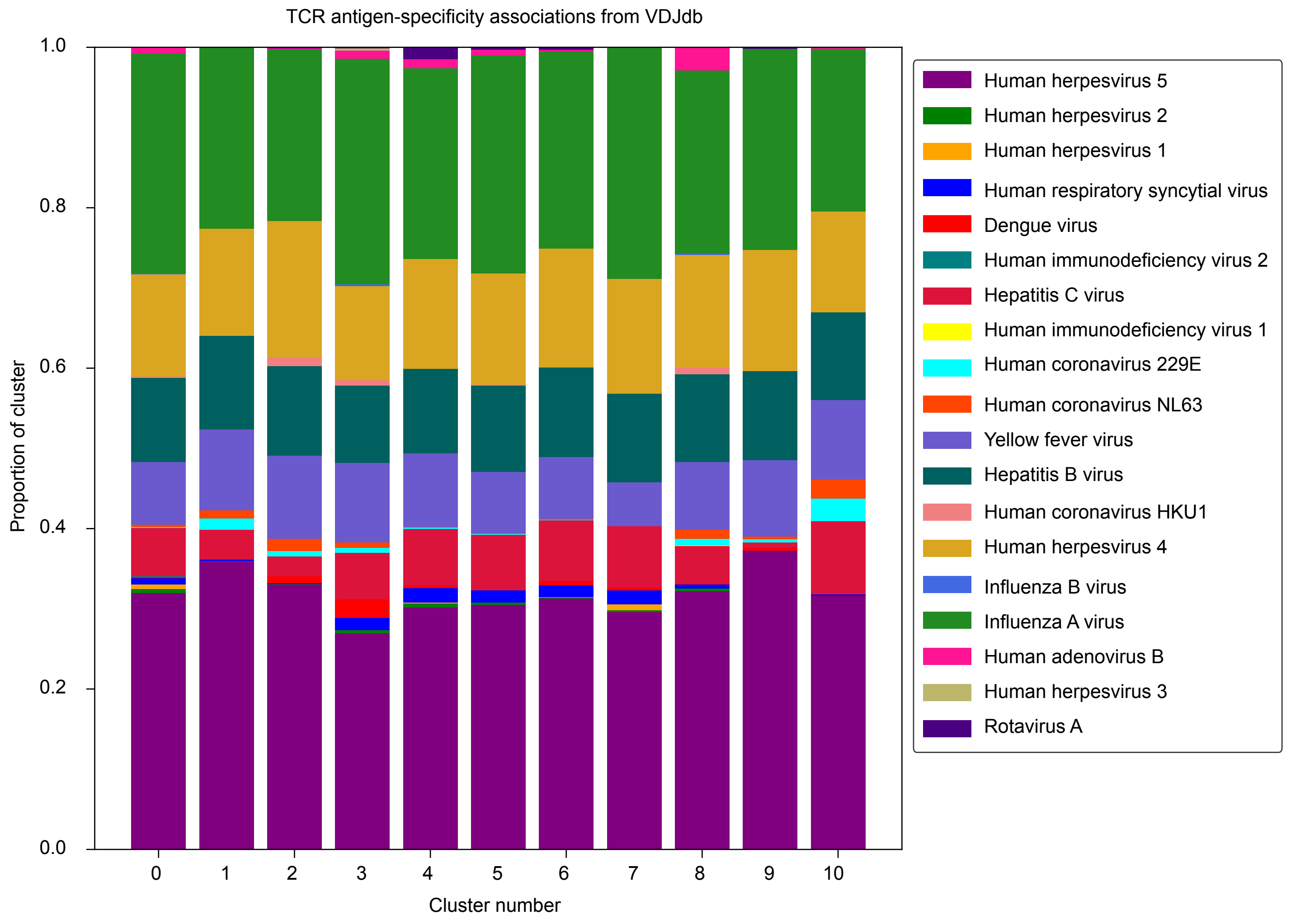

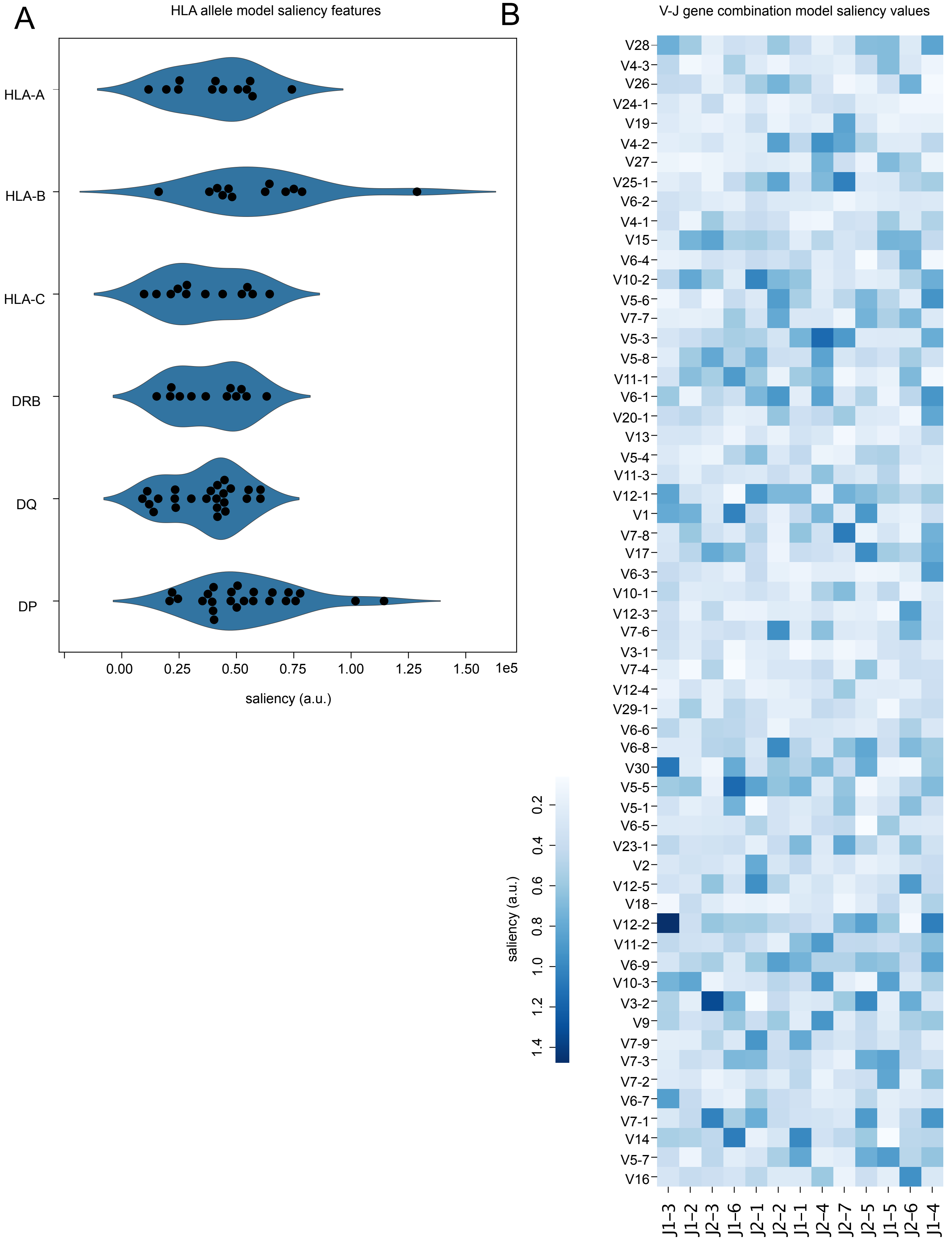
